# Composition of nasopharyngeal microbiota in individuals with SARS-CoV-2 infection across three COVID-19 waves in India

**DOI:** 10.1101/2023.01.02.522449

**Authors:** Tungadri Bose, Wasimuddin, Varnali Acharya, Nishal Kumar Pinna, Harrisham Kaur, Manish Ranjan, SaiKrishna Jandhyala, Tulasi Nagabandi, Binuja Varma, Karthik Bharadwaj Tallapaka, Divya Tej Sowpati, Mohammed Monzoorul Haque, Anirban Dutta, Archana Bharadwaj Siva, Sharmila S. Mande

## Abstract

Multiple variants of the SARS-CoV-2 virus have been plaguing the world through successive waves of infection over the past three years. Studies by independent research groups across geographies have shown that the microbiome composition in COVID-19 patients (CP) differ from that of healthy individuals (CN). However, such observations were based on limited-sized sample-sets collected primarily from the early days of the pandemic. Here, we study the nasopharyngeal microbiota in COVID-19 patients, wherein the samples have been collected across the three COVID-19 waves witnessed in India, which were driven by different variants of concern. We also present the variations in microbiota of symptomatic vs asymptomatic COVID-19 patients. The nasopharyngeal swabs were collected from 589 subjects providing samples for diagnostics purposes at Centre for Cellular and Molecular Biology (CSIR-CCMB), Hyderabad, India. CP showed a marked shift in the microbial diversity and composition compared to CN, in a wave-dependent manner. Rickettsiaceae was the only family that was noted to be consistently depleted in CP samples across the waves. The genera *Staphylococcus*, *Anhydrobacter*, *Thermus*, and *Aerococcus* were observed to be highly abundant in the symptomatic CP patients when compared to the asymptomatic group. In general, we observed a decrease in the burden of opportunistic pathogens in the host microbiota during the later waves of infection. To our knowledge, this is the first longitudinal study which was designed to understand the relation between the evolving nature of the virus and the changes in the human nasopharyngeal microbiota. Such studies not only pave way for better understanding of the disease pathophysiology but also help gather preliminary evidence on whether interventions to the host microbiota can help in better protection or faster recovery.

## Introduction

SARS-CoV-2 is the causative agent for the pandemic Coronavirus Disease 2019 where there have been 764,474,387 confirmed cases of COVID-19, including 6,915,286 deaths as of 26^th^ April 2023 (<u>WHO Coronavirus (COVID-19) Dashboard | WHO Coronavirus (COVID-19) Dashboard With Vaccination Data).</u> Since its onset, multiple variants of the virus have been reported, some of which have been categorized as variants of concern (VOC) viz. Alpha, Beta, Gamma, Delta, and Omicron [1]. While certain factors such as comorbidities, gender and age influence the onset of the disease as well as the severity of symptoms [2], the effect of the SARS-CoV-2 variants on the severity of COVID-19 disease is also speculated. Differences in transmissibility and entry pathways of the variants [3], [4] could have potentially given rise to the spectrum of symptoms that have been observed in the affected patients across waves [5], [6], [4]. Most patients showed signs of acute respiratory distress syndrome and typical symptoms such as fever, dry cough and tiredness (<u>Coronavirus disease (COVID-19) [who.int</u>] while other groups suffered from pain, anosmia, nasal congestion, sore throat and diarrhea [7]. Moreover, a majority of the population was asymptomatic of the viral infection thus, acting as a hidden carrier [8].

Human microbiota plays a significant role in modulating the host health by forging the immune system and it is also well known that the dysbiosis of the same has implications for diseases [9], [10], [11]. Therefore, association of the resident microbiota (or a dysbiosis thereof) with the symptoms and severity of the COVID-19 disease becomes worth exploring in context of different variants of concern. While some prominent studies in this direction have focused on gut microbiome dysbiosis in the COVID-19 infected patients [12], [13], [14]; recent reports also discuss the dynamics and the alterations of the nasopharyngeal microbiome [15], [16], [17]. However, these studies have mostly been restricted to the early waves of the pandemic, and do not include microbiome profiling during subsequent waves, which were caused by distinct virus variants (Fig. 1) and wherein the disease presentations were also different (first wave fueled by variant A2, second wave by Delta variant and third wave by the Omicron variant). Considering the strong association between viral and bacterial co-infections and respiratory disease severity, understanding the nasopharyngeal niche microbiome composition in context of the virulence/infectivity of different SARS-CoV-2 variants becomes imperative.

**Figure 1:**
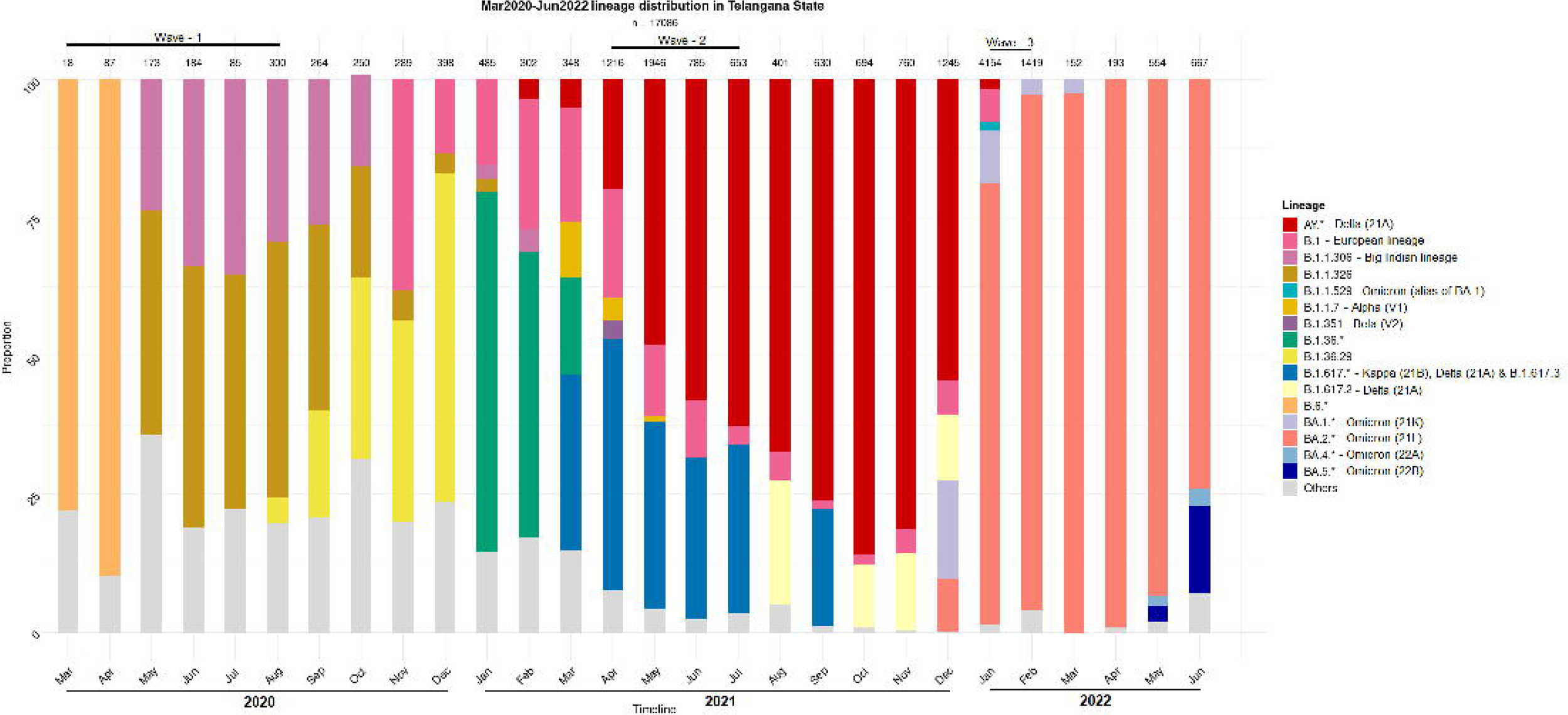
Distribution of the SARS-CoV-2 lineage among the samples collected from COVID-19 positive individuals. The nasopharyngeal samples were collected in the state of Telangana (India) between March 2020 and June 2022.

SARS-CoV-2 primarily enters the human body through ACE-2 and TMPRSS2 receptors present on the alveolar epithelial cells, and nasal epithelial cells of the nasopharyngeal tract and gradually moves towards the lungs [18], [19]. The delta and omicron variants have been observed to utilize different entry pathways for infection [3], and it is likely that during this migration, the resident microbiota can get altered and any inclusion of the pathobionts may aggravate the disease outcome. Studies by independent researchers have reported that nasopharyngeal microbiota during COVID-19 is characterized by a general decrease in the abundance of commensal organisms along with an increase in the abundance of opportunistic pathogens [20], [21], [22]. However, the individual organisms, both commensal as well as opportunistic, reported in each study seemed to vary. While this difference could be attributed to the choice of experimental protocols; geographical/ethnic differences between the subjects involved in each of the studies as well as the relatively lower number of samples analyzed in these studies, are also expected to play a major role in the study outcome. For instance, the opportunistic pathogens identified to be associated with the COVID-19 microbiota in an Indian study [23] differed considerably from those reported in studies conducted in other geographies [17], [21], [22]. Consequently, it is crucial to understand the nasopharyngeal microbiota signature of COVID-19 patients in India in the context of variants as well as waves through study of a larger cohort.

With the above objectives in mind, human nasopharyngeal microbiota was profiled from Indian subjects. The samples were provided for diagnostics and sequencing of SARS-CoV-2 at Centre for Cellular and Molecular Biology (CSIR-CCMB), Hyderabad, India. The samples collected from the subjects/ patients were grouped into four categories based on COVID-19 infection status (positive or negative) as well as symptom presentation (symptomatic or asymptomatic), and further processed for high throughput 16S rRNA gene amplicon-based sequencing for microbiome profiling. In addition to uncovering the variations between microbial signatures of COVID-19 positive patients (CP) versus COVID-19 negative individuals (CN), the study also reports how the nasopharyngeal microbiota changed during the pandemic over the three waves in India.

## Results

### Data overview

Nasopharyngeal swabs samples analyzed from 589 individuals between March 2020 – February 2022 (i.e., spanning three ‘waves of COVID-19 infection’ in India) were subjected to amplicon sequencing of the bacterial V4 hypervariable region of 16S rRNA gene on the Illumina MiSeq platform (details in Methods section). The category and wave-wise distribution of the 589 samples viz., symptomatic COVID-19 positives (sCP), asymptomatic COVID-19 positives (aCP), asymptomatic COVID-19 negatives or apparent healthy controls (aCN) and COVID-19 negatives exhibiting symptoms (sCN) [i.e., due to other respiratory/other infections] has been provided in Table 1. A total of 8645 amplicon sequence variants (ASVs) were identified in the microbiome sequence data (Supplementary Fig. 1) using the DADA2 package. Preliminary taxonomic analysis indicated the dominance of the phylum Firmicutes, Proteobacteria, and Actinobacteriota across all samples (cumulative abundance over 75%) (Supplementary Fig. 2a-b). At the family level, Staphylococcaceae and Corynebacteriaceae were found to be the most abundant taxa (Supplementary Fig. 2c-d).

**Table 1:** Statistics on the number of samples analyzed in the study. Categorization of 589 analyzed samples into sub-groups and their distribution across the three COVID-19 waves have been provided.

|  |  |  |  |  |  |
| --- | --- | --- | --- | --- | --- |
|  |  | Covid Positives (CP) |  | Covid Negatives (CN) |  |
|  | Category | with<br>symptoms (sCP) | without<br>symptoms (aCP) | with<br>symptoms (sCN) | asymptomatic<br>apparently<br>healthy<br>controls (aCN) |
|  | Total | 147 | 157 | 81 | 204 |
|  | Wave1 | 19 | 70 | 12 | 80 |
|  | Wave2 | 81 | 39 | 60 | 37 |
|  | Wave3 | 47 | 48 | 9 | 87 |

### Reduced microbial diversity during SARS-CoV-2 infection

To test whether COVID-19 status has any impact on the nasopharyngeal microbiome, we calculated the microbial alpha diversity between COVID-19 positive (CP) and COVID-19 negative samples. Using model selection based on the information-theoretic (IT) approach (Model Selection and Multimodel Inference: A Practical Information-Theoretic Approach | SpringerLink), [24] we found strong support for an effect of COVID-19 status, wave and combined effect of COVID-19 status & wave (COVID-19*Wave) on observed ASVs (ΔAIC_C_ = 38.47, R^2^_GLMM(m)_ = 0.312, R^2^_GLMM(c)_ = 0.320, Fig. 2a-b). Similarly, Shannon diversity, was observed to be influenced by COVID-19 status and Wave, individually, but not by COVID-19 status & wave together (ΔAIC_C_ = 22.85, R^2^_GLMM(m)_ = 0.277, R^2^_GLMM(c)_ = 0.312, Fig. 2c-d). For both alpha diversity indices, in general, lower diversity was observed in the COVID-19 positive (CP) samples as compared to the COVID-19 negative (CN) category (Fig. 2a-d). The effect of other variables included in the models such as age group, symptoms, gender, Ct-value were poorly supported by AIC_C_ model comparison for both alpha diversity indices (data not shown). Additionally, only the third wave samples showed a significantly lower diversity for observed ASVs in the CP category as compared to CN (p<0.05, Tukey’s HSD) which was reflected in Shannon diversity as well (all p>0.05). Overall, the results show a strong effect of COVID-19 infection on microbial alpha diversity indices, with the observed ASVs also showing an effect in a wave-dependent manner. Furthermore, the results suggest that this effect may be driven more by rare taxa than abundant ones, since the effect was less pronounced when accounting for abundance (i.e., Shannon diversity).

**Figure 2:**
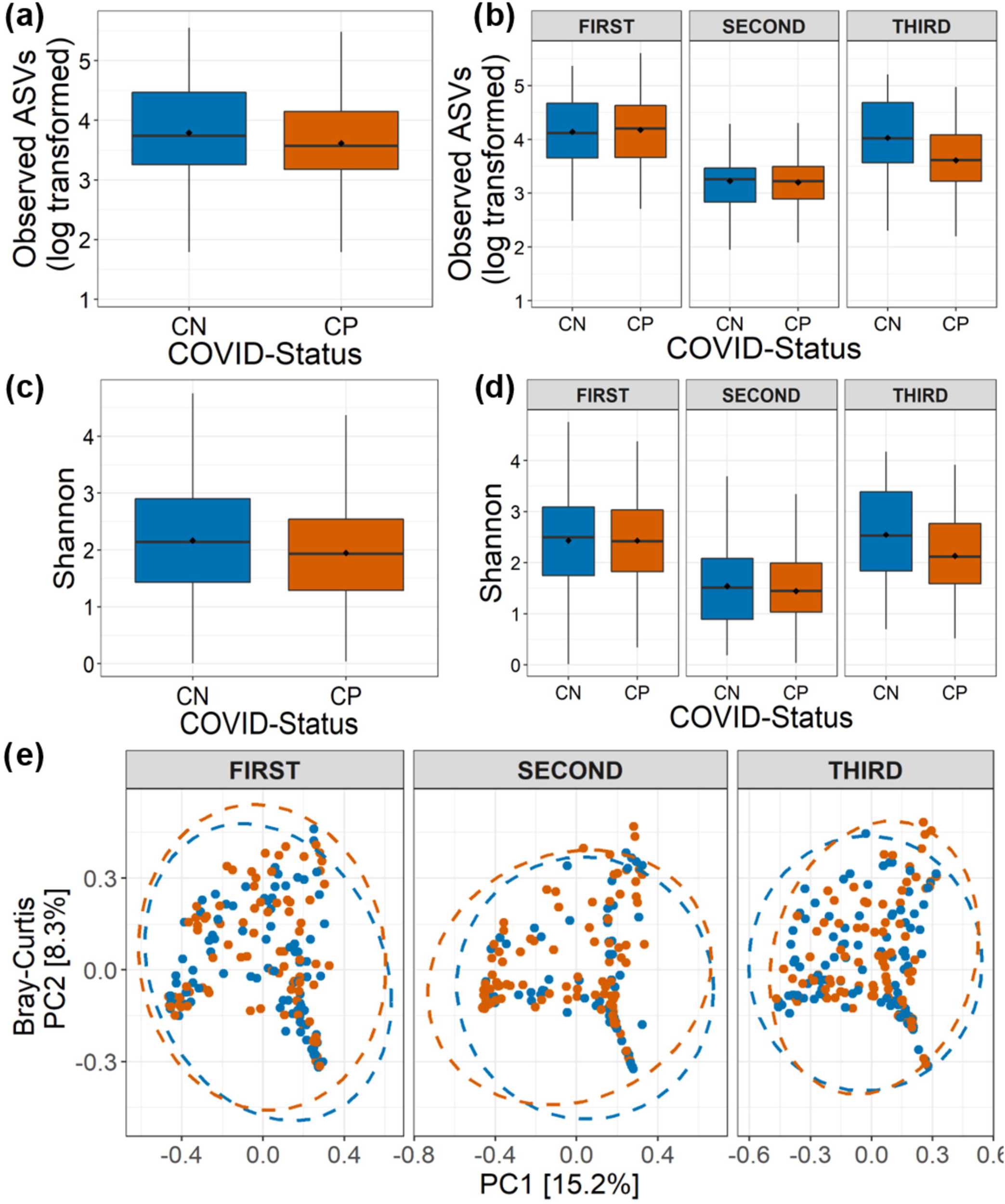
Alpha and beta diversity of the nasopharyngeal microbiota in COVID-19 positive (CP) and COVID-19 negative (CN) samples. Number of observed amplicon sequence variants (ASVs) (a) in all the analyzed samples (overall), and (b) in the three COVID-19 waves are shown. Shannon diversity in term of ASVs (c) in all the analyzed samples, and (d) in the three COVID-19 waves are depicted. (e) Beta diversity of microbiota assessed using principal component analysis (Bray-Curtis distance) represented along first two principal components for the three COVID-19 waves are presented.

### Community composition perturbed in COVID-19 nasopharyngeal microbiome

We also assessed the influence of COVID-19 status on the nasopharyngeal microbial community composition. While sequencing run (Jaccard: R^2^ = 0.010, p = 0.002; Bray-Curtis: R^2^ = 0.011, p = 0.004) and sequencing depth (Jaccard: R^2^ = 0.011, p = 0.001; Bray-Curtis: R^2^ = 0.016, p = 0.001) did impact the community compositions, controlling for these factors, the PERMANOVA models indicated a strong support for, the influence of COVID-19 status (Jaccard: R^2^ = 0.003, p = 0.006; Bray-Curtis: R^2^ = 0.003, p = 0.007), wave (Jaccard: R^2^ = 0.026, p = 0.001; Bray-Curtis: R^2^ = 0.037, p = 0.001) and the interaction between them (COVID-19*Wave) (Jaccard: R^2^ = 0.005, p = 0.001; Bray-Curtis: R^2^ = 0.006, p = 0.002) on the nasopharyngeal microbial beta diversity estimates (Fig. 2e). Additionally, Ct-value (Jaccard: R^2^ = 0.003, p = 0.005; Bray-Curtis: R^2^ = 0.003, p = 0.004) and symptoms (Jaccard: R^2^ = 0.002, p = 0.034; Bray-Curtis: R^2^ = 0.002, p = 0.017) also influenced the community composition. However, no effect of age group or gender (all p>0.05) on the nasopharyngeal microbial community composition was observed. PERMDISP (https://github.com/vegandevs/vegan) tests showed true shift in microbial community composition and no dispersion effect (all p>0.05) (Fig. 2e).

### Microbes associated with disease status across different COVID-19 waves

Given the observed variation of microbial taxonomic diversity across three COVID-19 waves, negative binomial Wald tests were performed using the DESeq2 package with the intent to identify the taxa that are differentially abundant between CP and CN groups corresponding to different waves. While there were no significant differences (BH corrected p<0.05) in the first wave samples, there was a marked compositional variation in the CP microbiota of the second and third wave. Cyanobacteria and Firmicutes were the dominant phyla, with Planctomycetota, Deinococcota, and Bdellovibrionota having a significantly lower abundance in the CP microbiota of the second wave samples (Supplementary Fig. 3). Similarly, third wave samples showed a significantly enriched proportion of Proteobacteria in CP compared to CN.

At the family level, amongst the taxonomic groups discriminating between CP and CN in each of the three waves, Rickettsiaceae was found to be depleted in all the three waves as seen in Fig 3. A small number of taxa showed significant differences between CP and CN microbiota in the first wave (enriched abundance of Hymenobacteraceae and Spirosomaceae with a decrease in the abundance of Thermaceae and Rickettsiaceae in CP). Additionally, the CP group of second wave samples were enriched with five bacterial families (Rhizobiales.Incertae.Sedis, Hydrogenophillaceae, Alteromonadaceae, Hymenobacteraceae and Brevibacteriacceae) while two of them (Devosiaceae and Thermaceae) had a lower abundance. In the case of the third wave, three families (viz. Carnobacteriaceae, Burkholderiaceae, and Corynebacteriaceae) had a higher abundance whereas seven families (viz. Aeromonadaceae, Rickettsiaceae, Alteromonadaceae, Pasteurellaceae, Spirosomaceae, Aerococcaceae, Fusobacteriaceae) exhibited a lower abundance in the CP group. Furthermore, at the genera and ASV levels, the genera *Aliterella* and ASVs: ASV36 (family Rickettsiaceae), ASV161 (*Gemella*), ASV195 (*Deinococcus*), and ASV431 (*Elizabethkingia*) were noted to demonstrate a significant decrease in abundance in the CP group across all the three infection waves (Supplementary Table 1).

**Figure 3:**
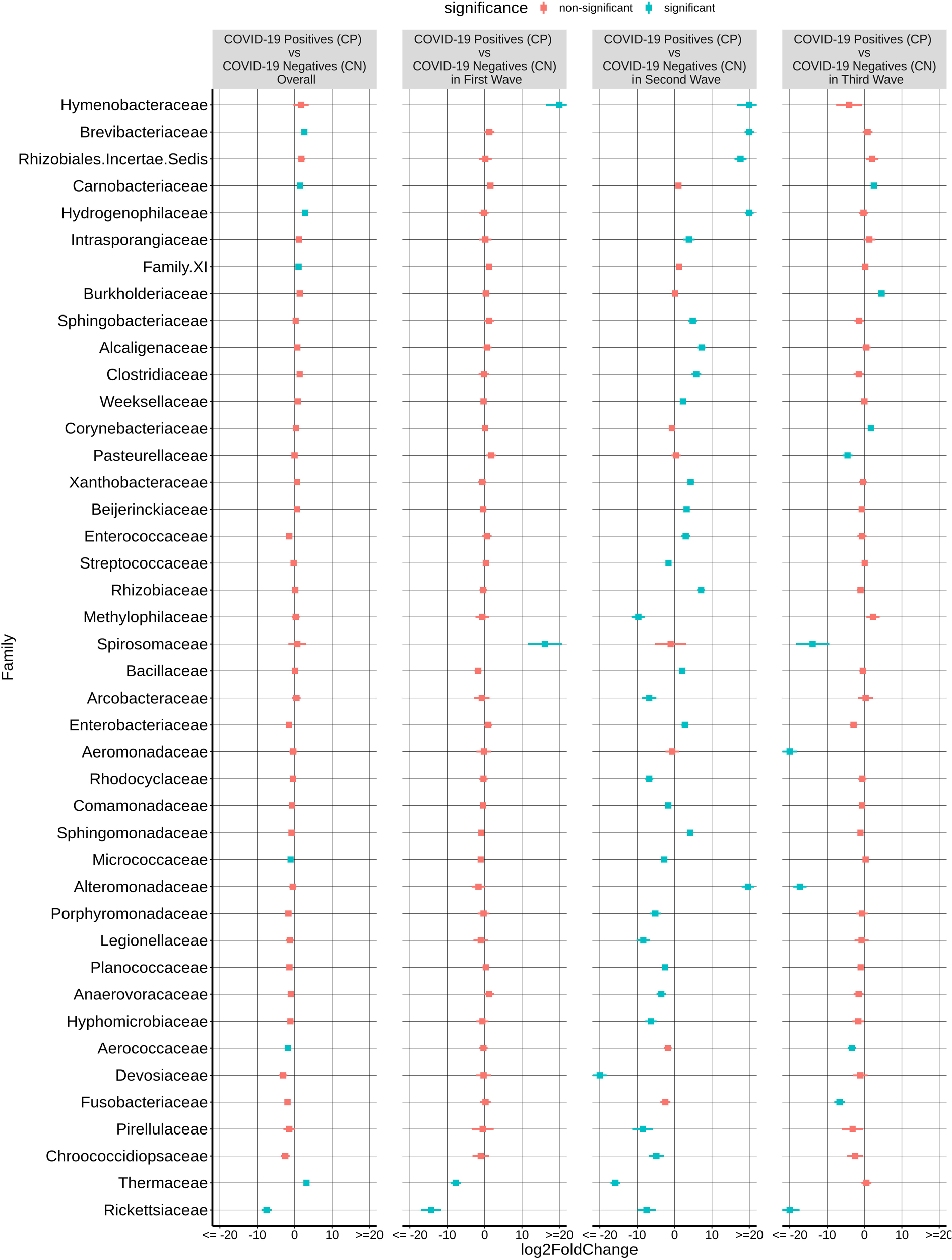
Differential abundance of bacterial families between the COVID-19 positive (CP) and COVID-19 negative (CN) samples. The log2fold change in the mean abundance (along with whiskers representing standard errors) of a bacterial family in CP with respect to CN is depicted in (a) all the analyzed samples (overall) as well as in (b-d) each of the three COVID-19 waves. Significantly different abundance (q-value < 0.05) is indicated with blue colour.

### Microbes associated with inter-wave variations in CP and CN groups

Most CP cases particularly during the second and the third Covid-19 wave in India were dominated by one of the viral variants (Fig. 1). Supplementary Fig. 4 shows the effect of different waves (and thus viral variant) on the bacterial taxonomic proportions observed in CN and CP groups of samples in terms of fold changes at the phylum level. As expected, based on the observations from the alpha and beta diversity tests, the microbiome of the second wave samples was distinct from the other two. When compared to the first wave, the second wave samples were found to be significantly enriched with the phyla Proteobacteria, Campylobacterota and Firmicutes, in both CP and CN groups. In contrast, Cyanobacteria, Planctomycetota, Bdellovibrionota and Fusobacteriota was depleted in both the CP and CN samples from the second wave when compared to those from the first wave. When the third wave samples were compared with those from the second wave, the phyla Cyanobacteria, Plantomycetota and Bdellovibrionota were significantly enriched in the third wave in both CP and CN samples. On the other hand, only Firmicutes were observed to be significantly depleted in the CP and CN samples of the third wave when compared to the second wave. The first wave and the third wave samples were observed to be more similar to each other. The only significant differences pertained to enrichment of Deinococcota and depletion of Bacteriodota in the CP and CN samples of the third wave compared to the first wave samples. It was intriguing to note that the phylum Deinococcota showed a gradual (and in most cases significant) increasing trend over the three waves in both CP and CN groups.

Additional results depicting the wave effect on microbiota at genera and ASV levels are provided in Table 2. Among the CP group, the genera *Abiotrophia*, *Streptococcus*, *Rheinheimera* along with *X.Eubacterium*..brachy.group and *Escherichia.Shigella* were found to follow a constant decrease in abundance between the corresponding waves (i.e., these genera were most abundant in the first wave CP samples and their abundances were least in the third wave CP samples). In the CN group, *Schlegelella* had a significantly increasing trend, whereas Acinetobacter and *Megasphaera* showed a significant decreasing pattern. Supplementary Table 2 shows a list of the ASVs and genera which followed a consistent pattern of significant increase or decrease in abundance in the CP and CN groups.

**Table 2:**
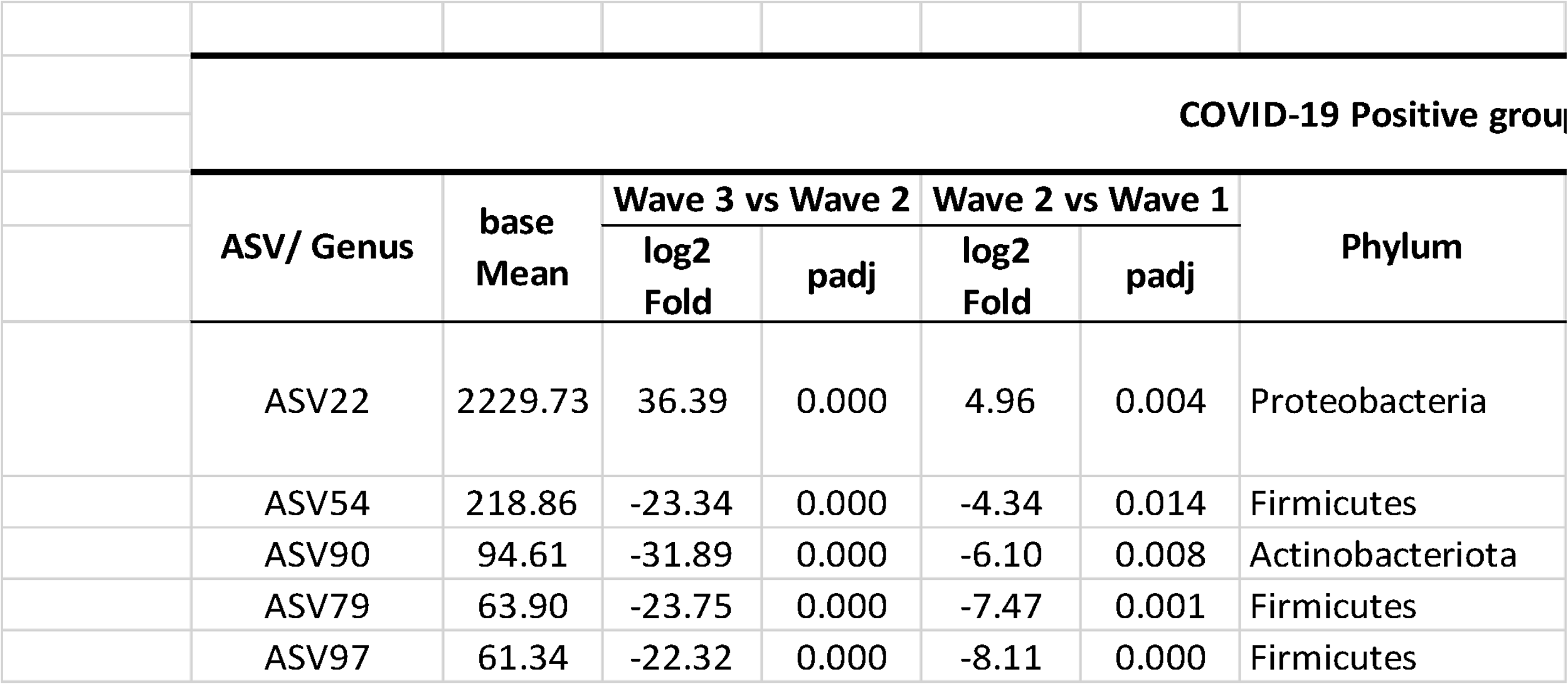
List of the amplicon sequence variants (ASVs) and genera demonstrating a consistent increase or decrease in abundance across the three COVID-19 waves. Significant changes having an adjusted p-value < 0.05 have been reported.

### Microbial taxa associated with symptomatic and asymptomatic COVID-19 subjects

Supplementary Fig. 5 shows the overall bacterial taxonomic proportions observed in CN and CP groups of samples categorized based on symptom in terms of fold changes at the phylum levels (abundance plots are seen in Supplementary Fig. 2b & d). Within COVID-19 positive (CP) group, symptomatics (sCP) had a significant overabundance of the phyla Camphylobacterota, Patescibacteria and Firmicutes, while showing a depletion in the abundance of the phyla Fusobacteria and Proteobacteria compared to the asymptomatics (aCP).

At the family level, sCP was observed to have a significantly higher abundance of Rickettsiaceae, Aerococcaceae and Thermaceae among others as well as a significantly lower abundance of Leptotrichiaceae, Weeksellaceae, Deinococcaceae, Sphingomonadaceae and Xanthobacteraceae (the latter two belonging to phylum Proteobacteria) when compared to aCP (Fig. 4). The severity of COVID-19 infection (i.e., sCP vs aCP) appeared to be linked to a higher abundance of the family Saccharimonadaceae, and Chitinophagaceae, and a lower abundance of the family Sphingomonadaceae, and Weeksellaceae.

**Figure 4:**
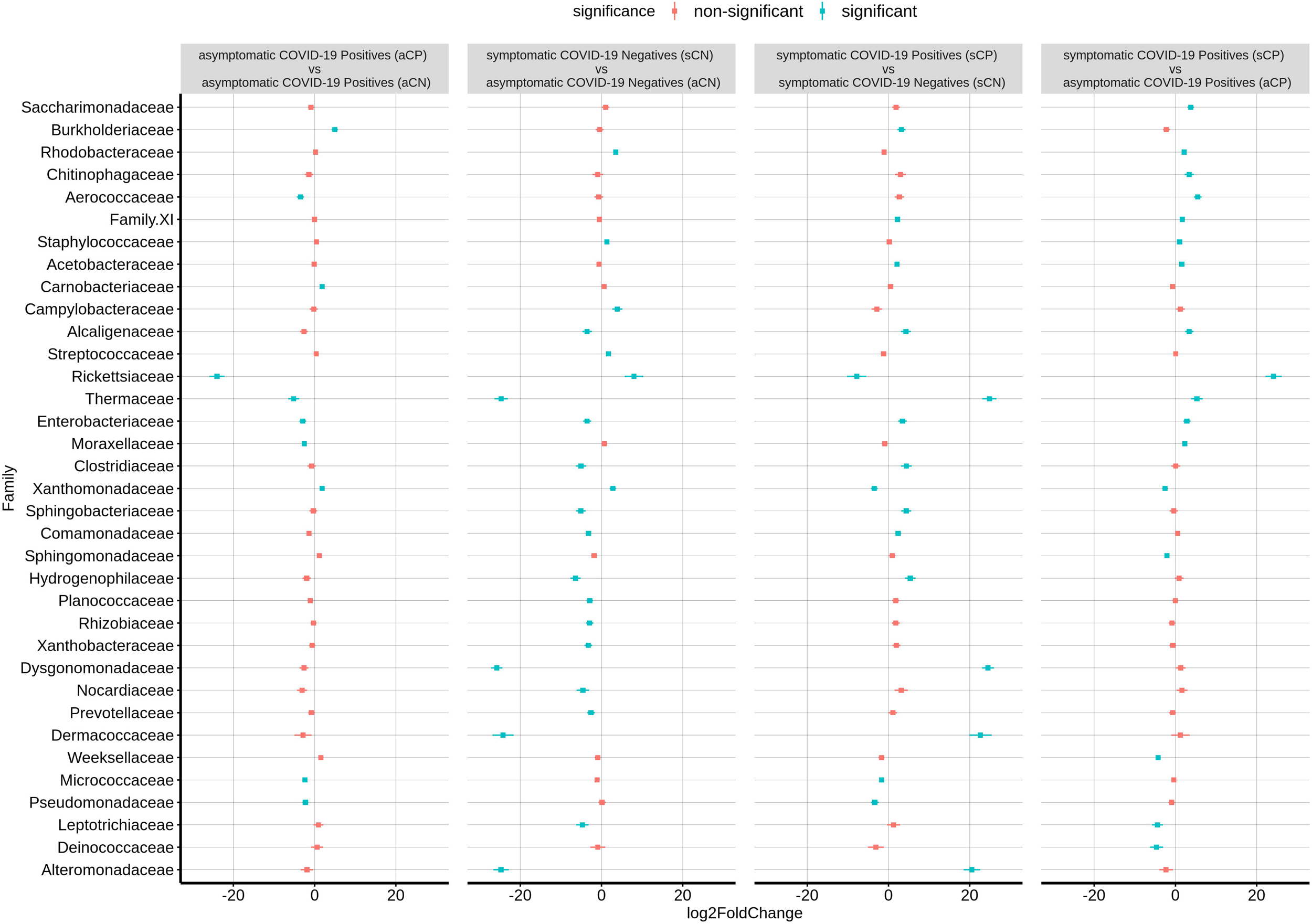
Differential abundance of bacterial families between the four sub-group of samples - asymptomatic COVID-19 positive (aCP), symptomatic COVID-19 positive (sCP), apparently healthy controls who are COVID-19 negatives as well as asymptomatic (aCN), and COVID-19 negatives exhibiting symptoms (sCN). The log2fold change in the mean abundance (along with whiskers representing standard errors) of a bacterial family in (a) aCP with respect to aCN, (b) sCN with respect to aCN, (c) sCP with respect to sCN, and (d) sCP with respect to aCP is depicted. Significantly different abundance (q-value < 0.05) is indicated with blue colour.

The list of the significant changes in abundance of microbes at a genus level between sCP and aCP have been provided in Table 3. *Thermus, Aerococcus*, *Enhydrobacter, and Staphylococcus* were found to be the most significantly enriched genera in sCP when compared to aCP. ASV108 (*Thermus amyloliquefaciens*) was found to be enriched in sCP in comparison to both aCP (Supplementary Table 3) indicating that its levels in the nasopharyngeal microbiota might act as an early indicator of the severity of the disease outcome. In contrast, *Elizabethkingia, Leptotrichia,* and *Veillonella* were more abundant in aCP (w.r.t. sCP) (Table 3). Additional insights on how the microbiota profiles of the symptomatic and asymptomatic COVID-19 positive patients compare with that of COVID-19 negative subjects with or without symptoms have been provided in the Supplementary Results and Supplementary Table 3.

**Table 3:** List of the bacterial genera which showed significant changes in abundance between symptomatic COVID-19 positive (sCP) and asymptomatic COVID-19 positive (aCP) sub-groups. Significant changes having an adjusted p-value < 0.05 have been reported. Positive log2foldchange values indicate a higher abundance in sCP while negative log2foldchange values represented genera with higher abundance in aCP.

| Genus | base Mean | log2 Fold Change | padj | Phylum | Family |
| --- | --- | --- | --- | --- | --- |
| Elizabethkingia | 89.341 | -4.8943 | 0.0019 | Bacteroidota | Weeksellaceae |
| Leptotrichia | 49.941 | -4.8144 | 0.0085 | Fusobacteriota | Leptotrichiaceae |
| Veillonella | 245.31 | -2.6977 | 0.0145 | Firmicutes | Veillonellaceae |
| Staphylococcus | 54139 | 1.2499 | 0.003 | Firmicutes | Staphylococcaceae |
| Enhydrobacter | 161.48 | 2.8906 | 0.0411 | Proteobacteria | Moraxellaceae |
| Thermus | 26.63 | 4.7846 | 0.0145 | Deinococcota | Thermaceae |
| Aerococcus | 713.23 | 9.1933 | 3E-11 | Firmicutes | Aerococcaceae |

### Microbial cooccurance networks varied with COVID-19 status across the three waves

The microbial association network (Pearson correlation) corresponding to each of the three COVID-19 waves, CP, CN as well as those for symptomatic and asymptomatic conditions were constructed considering the taxonomic abundance data at the genera level. Network properties of each of these networks are provided in Table 4. In line with observations pertaining to diversity indices, minimal number of nodes (genera) and edges (interactions between nodes) were observed in the network corresponding to the second wave whereas the number of nodes and edges in the third wave network was observed to be considerably higher than that in the first two waves. The distribution of betweenness centralities of the nodes in the network seemed to indicate a considerable shift in the network architecture between the subsequent waves (Supplementary Fig. 6). As mentioned earlier, with respect to the first wave, we noted a loss of many nodes in the network corresponding to the second wave, including the genus *Actinomyces* which appeared to be a high betweenness node (degree = 14; betweenness =0.618; stress = 74) in the microbial network corresponding to the first wave (Supplementary Fig. 7). The change in network architecture from the first to the second wave was driven by the genera *Cnuella*, *Marmoricola*, *Paenibacillus*, *Peptostreptococcus*, and *Solobacterium* as indicated by their NESH scores (Supplementary Table 4). Although the abundance of most of these genera (except *Cnuella*) in the second wave was lower in both CP and CN samples w.r.t. first wave (Supplementary Table 2), the driver taxa were found to disrupt the sub-network of the first wave, thereby leading to an altered association among the genera in the network representing the second wave. *Cnuella* was higher in abundance only in the CP group, while its abundance decreased in the CN group. Notably, some organisms belonging to the genera *Paenibacillus*, *Peptostreptococcus*, and *Solobacterium* are known to be opportunistic pathogens [25], [26], [27]. Further, some of these driver taxonomic groups, viz., *Cnuella*, *Marmoricola*, and *Paenibacillus* lost their importance in the microbiome association network corresponding to the third wave (Supplementary Fig. 8). *Leptotrichia* and *Actinomyces* were noted to be taxonomic groups driving the changes during the third wave when compared to the second wave. While the abundance of *Actinomyces* decreased, the abundance of *Leptotrichia* was higher in the third wave w.r.t. the second wave (Supplementary Table 2). Further, *Leptotrichia* appeared as a node with high stress centrality (degree = 18; betweenness =0.317; stress = 236) in the third wave network.

**Table 4:** Properties of the microbial association networks analyzed in this study.

|  | Network | No of Nodes | No of Edges | Avg No of Neighbours | Characteristic Path Length | Clustering Coefficient |
| --- | --- | --- | --- | --- | --- | --- |
|  | Wave 1 | 85 | 278 | 5.5 | 1.808 | 0.774 |
|  | Wave 2 | 56 | 92 | 3.33 | 1.4 | 0.667 |
|  | Wave 3 | 92 | 486 | 4.7 | 2.284 | 0.586 |
|  | Symptomatic | 49 | 128 | 5.75 | 1.179 | 0.94 |
|  | Asymptomatic | 63 | 214 | 7.833 | 1.303 | 0.854 |
|  | Covid Positive | 69 | 214 | 2 | 3.091 | 0.091 |
|  | Covid Negative | 75 | 210 | 4.727 | 1.673 | 0.596 |

In case of CP and CN networks, the CP-network was characterized by longer path lengths (3.091) compared to the CN-network (1.673). The network density values indicate a denser CN-network compared to the CP-network. The number of nodes with high betweenness values were also higher in the CN network compared to the CP network. The interacting partners for the microbes were seen to considerably change from the CN to CP. This resulted in the generation of alternate microbial subnetworks between the two conditions which might have an implication of the behavior of the individual microbes (Fig. 5 a & b). Major changes were observed in clusters 2, 3 and 4 of the CN networks. Several members of cluster 2, such as *[Eubacterium] nodatum* group, *Atopobium*, and *Mogibacterium* were characterized by an increased NESH score in the CP network. Notably, the genera *Schlegelella* which was earlier found to consistently increase in abundance across the three waves of the COVID-19 pandemic in the CN samples (Table 2), witnessed a drop in its betweenness centrality measure in the CP network, thereby indicating at its strong association with the CN nasopharyngeal microbiota (Fig. 5). The insights on the shift in the microbial associations between the asymptomatic and symptomatic networks (Supplementary Fig. 9) have been provided as Supplementary Results.

**Figure 5:**
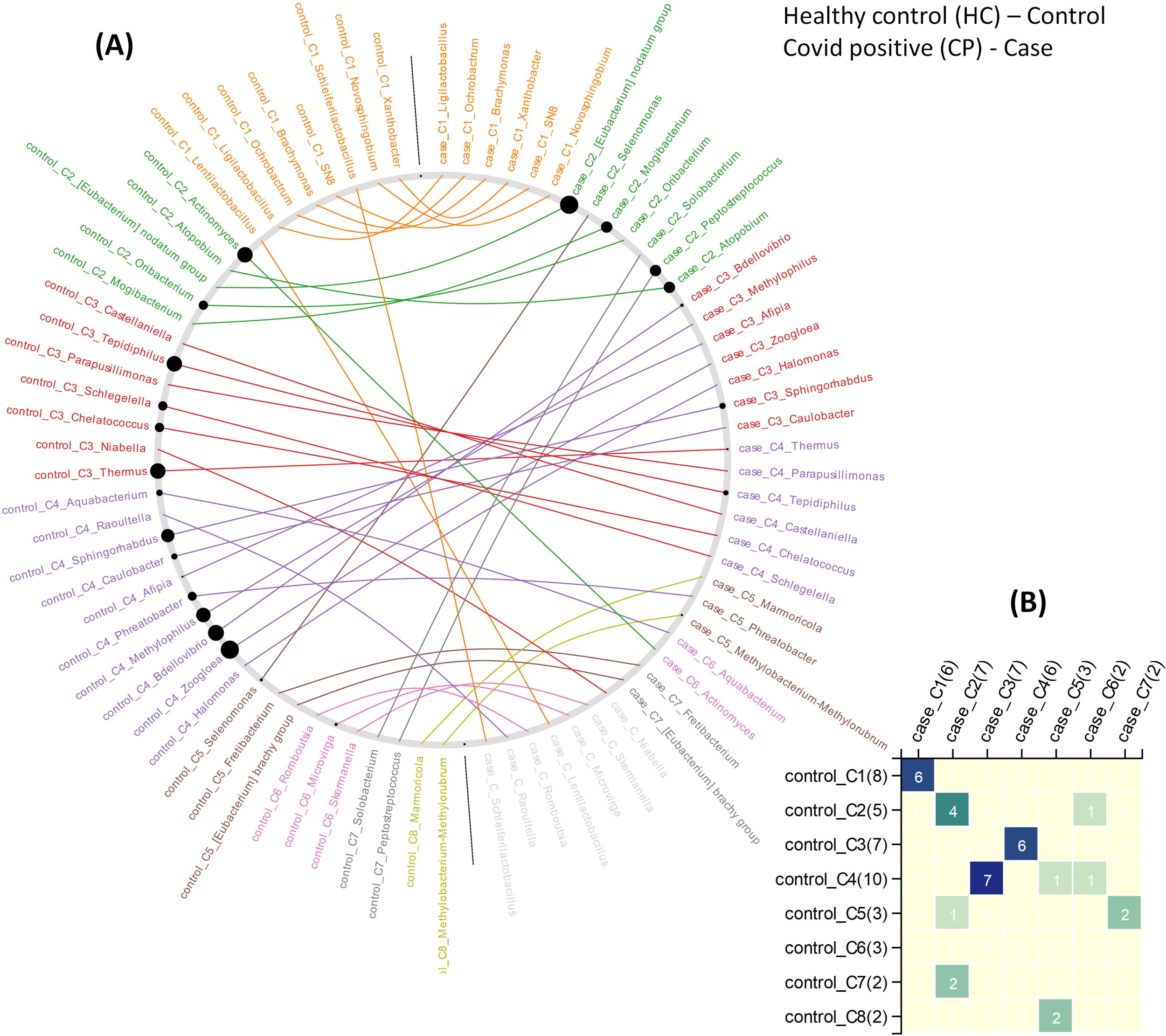
The changes in community structure (community shuffling) between the microbial association networks corresponding to the COVID-19 negative (CN) - ‘control’ and COVID-19 positive (CP) - ‘case’ networks. (a) Nodes belonging to the CN - ‘control’ and CP - ‘case’ networks are plotted along the left half and right half of the circular frame. Same node (microbe) in the two network is connected by an edge for easy viewing of the community shuffling. Node labels are coloured (at random) based on sub-network/ community affiliations. Grayed out node labels indicate that the node does not interact directly with the common sub-network. The node sizes are proportional to the betweenness centrality measure of the node in the corresponding network. (b) Heatmap representing the communities in the CN - ‘control’ network (along the vertical axis) and the CP - ‘case’ network (along the horizontal axis) with counts of the nodes (microbes) in each community in brackets. Numbers (and colour gradient) in the heatmap indicates how the nodes constituting any sub-network/ community of the CN - ‘control’ network are shared with the sub-networks/ communities of the CP - ‘case’ network, and vice versa.

It may be appreciated that the interactions between the different microbial groups in a microbial network, i.e., their symbiotic relationships would be driven by the functional potential of the interacting microbes. Therefore, the inferred functional potentials of the microbes associated with the microbiota samples were also analyzed. The results of this analysis (Supplementary Tables 5-8) and the subsequent insights from the analysis have been provided in the Supplementary Results section.

## Discussion

The present study aimed at profiling the nasopharyngeal microbiota of Indian subjects, across the three COVID-19 waves, to identify signatures, if any, which may be distinct between the microbiota sample collected from COVID-19-positive patients (CP) and negative (CN) individuals. It may be noted that the different infection waves were caused by different viral variants. In particular, the second and the third COVID-19 wave was predominated by the Delta (21A) and Omicron (21L) variants respectively (Fig. 1). Given that subtle variations exist in the pathophysiology of different viral variants [3] [28], it was anticipated that the associated host microbiota would also exhibit certain changes in each of the three waves. While the Supplementary Discussion sections put forth the more generic observations, the salient observations made in the context of COVID-19 disease are being discussed here.

Our findings indicate decreased microbial diversity associated with COVID-19 infections. Earlier studies too have shown that the diversity of the microbes constituting the nasopharyngeal microbiota decreased in patients with confirmed COVID-19 infections [17], [29], [23], barring a few exceptions [30]. A study by Gupta A et al. on a relatively smaller cohort in India had further identified host type (age and gender) and patient condition (symptomatic vs. asymptomatic) as potential factors affecting the enrichment of specific bacterial communities in upper respiratory tract [23]. In our study, while the condition of the patients (i.e., symptomatic vs. asymptomatic) was found to be associated with bacterial diversity alone; while age and gender did not seem to significantly influence the same (Supplementary Fig. 5).

It may be noted that the depletion of Rickettsiaceae in CP samples was the only consistent pattern that was observed across the three COVID-19 waves at a family level. At an ASV level, this observation was supported by a significant drop in the abundance of ASV36 in CP w.r.t. CN (Table 2). Notably, while the abundance of Rickettsiaceae decreased during COVID-19 infections, it has been reported to increase in other respiratory infections [31]. Thus, it appears to be a distinguishing feature between COVID-19 and other respiratory infections. To our knowledge, this is the first report on the (negative) association of the abundance of Rickettsiaceae with CP. Further studies into this aspect will be required to see if the presence of Rickettsiaceae in the nasopharyngeal microbiota could provide any advantage in combating the onset/progression of a COVID-19 infection.

Other consistent patterns across the three COVID-19 waves included depletion of the genera *Aliterella* as well as certain ASVs from the genera *Deinococcus* (ASV195) and *Elizabethkingia* (ASV431) in CP. This is consistent with a previous study involving SARS-CoV-2 infected patients which have also shown the absence of microbes belonging to the phylum Deinococcus-Thermus in patients admitted to ICU [32]. It was intriguing to note that despite multiple reports on the risk and incidences of septicemia caused by *Elizabethkingia* in COVID-19 patients [33] [34] the abundance of *Elizabethkingia* was found to be lower in CP compared to CN. While the causation of septicemia in most case has been attributed to *E. meningoseptica* [33] [34] the ASV corresponding to *Elizabethkingia* (ASV431) identified in the current study matched closer to *E. anopheles* (with 100% query coverage and 99.6% sequence identity) when compared to *E. meningoseptica* (having 100% query coverage and 98.42% sequence identity). Given an earlier report on the contamination of throat swab collection kits with *E. anopheles* we presume that the observed enrichment of *Elizabethkingia* in CN could also be an artifact.

In contrast, *Gemella massiliensis* (ASV161) was enriched in CP across all the three waves. Interestingly there have been contradictory reports regarding the association of *Gemella* with COVID-19 infection. While studies conducted in China had reported a depletion of *Gemella* species like *G. morbillorum* and *G. haemolysans* in swab samples collected from the pharynx of COVID-19 patients [22] a study involving subjects in India had shown an increment in the abundance of *Gemella* to be associated with CP [23]. It is therefore likely that the association of *Gemella* with COVID-19 infection might vary across geographies. The observation pertaining to a lower abundance of genera *Aliterella* in CP across all the three COVID-19 waves was also noteworthy. While none of the previous literature on association of microbiota with COVID-19 reported a similar observation, a few works exploring the antiviral properties of Cyanobacteria drew our attention [35], [36]. Cyanobacteria such as *Aliterella* are a rich source of bioactive compounds and are likely to possess antiviral properties which supports their higher abundance in the CN group.

From the perspective of the potential role of microbiota in the manifestation of COVID-19 infection, our study reports the enrichment of IgA-specific metallo-endopeptidase enzyme (EC:3.4.24.13) in CP samples (Supplementary Table 6 - overall). This metal-dependent enzyme cleaves the Pro-Thr bond in the hinge region of the heavy chain in immunoglobulin A (IgA) and is known to be encoded by pathogenic bacterial groups like *Streptococcus* [37], [38]. Given the role of IgA in mucosal immunity [39] and the reports on the association of IgA with the criticality of COVID-19 disease [40], it appears that CP patients are indeed vulnerable to secondary bacterial infections.

Given the observed variations in the nasopharyngeal microbiota across the COVID-19 waves (Fig. 2, Fig. 3 and Supplementary Fig. 1-3), we were interested to find which of the changes in the microbiota followed similar trends in both the CP and CN groups. The microbiota abundance signature (at a phylum level) of the first and the third wave seemed to be similar with each other as compared to that of the second wave (Supplementary Fig. 2 & 3). Across both the CP and CN groups, the abundance of the phylum Deinococcota was found to gradually increase during the course of the pandemic (Supplementary Fig. 3). Although this increase in abundance of Deinococcota was not always statistically significant, the trend was very clear. A number of ASVs (and genera) were also observed to follow a consistent up/ down trend in either CP or CN group across the three waves (Table 2). Most notably, in the CP group, majority of the ASVs which demonstrated a downtrend during the course of the pandemic represented opportunistic pathogens like *Veillonella parvula*, *Rothia mucilaginosa*, *Streptococcus anginosus*, *Acinetobacter baumannii*, etc. Since, previous studies have reported an increase in the abundance of opportunistic pathogens in the microbiota of COVID-19 patients compared to healthy individuals [22], [16]; this observation might seem counter intuitive. It may be noted that most studies [22], [16], [23] reporting a higher abundance of opportunistic pathogens in microbiota samples from COVID-19 patients were based on data collected during the early part of the pandemic (samples collected in 2020). It is likely that during early days of the pandemic, the human immune system fighting the SARS-CoV-2 virus resulted in an immunocompromised state that could be exploited by the opportunistic pathogens to form secondary infections.

The findings by Devi *et al.* [41] regarding the higher transcript abundance of *Achromobacter xylosoxidans* and *Bacillus cereus* in cases of COVID-19 associated mortality, and *Leptotrichia buccalis* in the severe COVID-19 cases highlights the role of co-infecting microbes in the severity and outcome of COVID-19. Thus, despite the observations made in our study that implicated the perturbation of nasopharyngeal microbiota as a result of the viral infection, it cannot be completely ruled out that these changes could be a potential factor aggravating the infection and disease outcome. However, during the later waves, once vaccines against the SARS-CoV-2 virus were introduced, as well as there were many more cases with prior exposure to the pathogen; it is likely that the immune system has become better equipped to deal with the challenges posed by the virus as well as to ward off secondary infections caused by opportunistic pathogens. The lower abundance of bacteria with pathogenic potential like *Streptococcus matis*, *Acinetobacter baumannii*, *Lachnoanaerobaculum orale*, and *Megasphaera micronuciformis* even in the CN group also supports the above notion (Table 2). Indeed, among the genera which were identified to drive the change in microbial network architecture from the first to the second wave (Supplementary Fig. 6), several were noted to be opportunistic pathogens (such as, *Paenibacillus*, *Peptostreptococcus*, and *Solobacterium*) [25], [26], [27] whose abundance decreased in both CP and CN samples from the second wave when compared to the first wave. In addition to vaccination, lower exposure to pollutants due to lockdown and usage of masks as well as practicing home remedies for improving respiratory health might have also led to the overall decrease in the proportion of opportunistic pathogens in the nasopharyngeal tract of Indians over the course of the pandemic. An exception was noted in the case of the genera *Leptotrichia* which appeared to drive the change in the microbial network architecture (and increased in abundance) from the second to the third wave (Supplementary Fig. 7). Though part of commensal respiratory microbiota, species of the genera *Leptotrichia* genera (eg. *L. buccalis*) possess pathogenic properties and have been shown to have a higher abundance of transcripts in severe COVID-19 patients [41]. In this context, it is also important to note that we did not find any bacterial function which was consistently enriched or depleted in the CP group (w.r.t. CN) across the three COVID-19 waves. As stated above, the changing nature of the underlying microbiota structure through the course of the pandemic due to vaccination, lockdown and/or other interventions might be a reason for this observation.

The wide variation in the symptoms in COVID-19 positive patients led us to investigate whether there were any microbial association with the severity of the infection, i.e., between the symptomatic and the asymptomatic COVID-19 positive sub-groups (sCP; n = 147 vs aCP; n = 157). Our findings (see Table 3) differed from a recently published study by Gupta *et al.* which also inspected the same phenomenon using a smaller cohort of 11 symptomatic and nine asymptomatic COVID-19 positive patients [23]. While this could be due to variations in data analyses protocols, the differences in the number of samples in the two studies might have also influenced the outcome. In this context, we found a set of five ASVs (Table 5) whose abundance seemed to be linked to symptoms in response to infections in the respiratory tract. A thorough investigation into their role in disease manifestation might reveal whether and how these organisms interact with the virus and the host immune system in driving the disease outcome. Details of all the ASVs identified in this study have been provided in Supplementary Table 4.

**Table 5:**
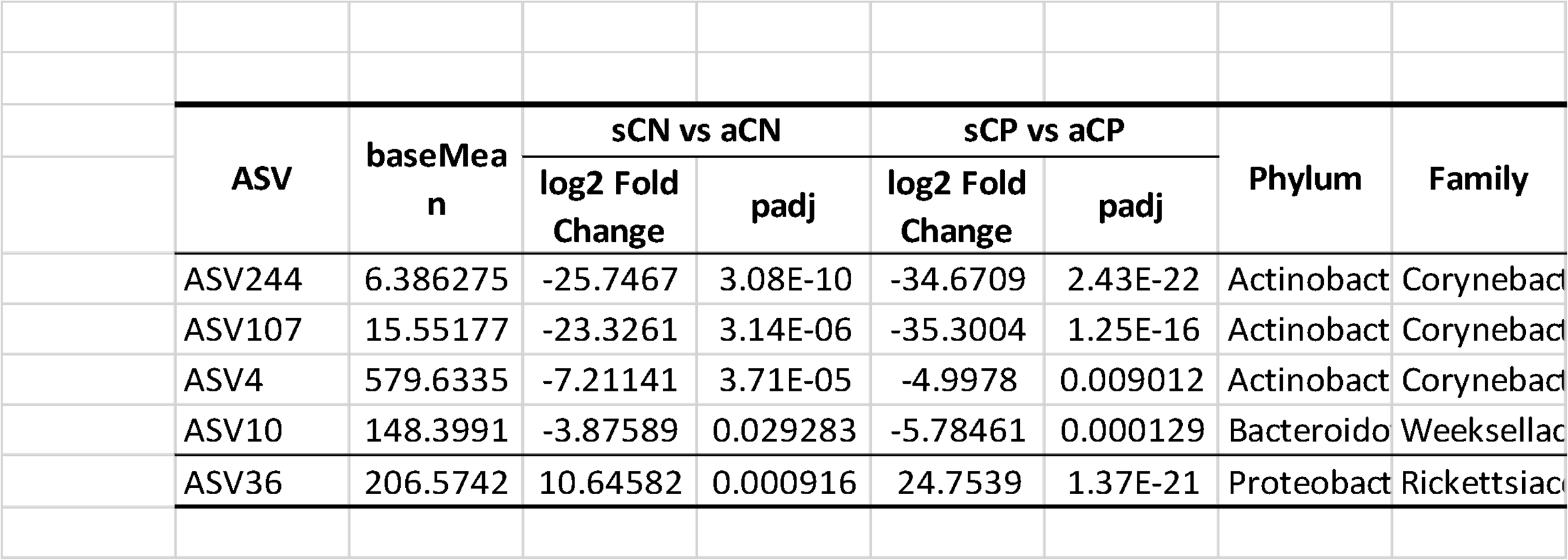
List of amplicon sequence variants (ASVs) whose abundances were associated with symptoms in response to infections in the respiratory tract. Significant changes having an adjusted p-value < 0.05 have been reported. Positive log2foldchange values indicate a higher abundance of the ASV in symptomatic samples, and vice versa.

In addition, we were also intrigued to find the enrichment of superpathway of mycolyl-arabinogalactan-peptidoglycan (mAGP) complex biosynthesis (PWY-6404) in the sCP when compared to sCN sub-group. Since mAGP complex is best known to be associated with the viability of *Mycobacterium tuberculosis*, we speculate that symptomatic COVID-19 positive (sCP) patients have a likelihood to activate latent tuberculosis infection. Some of the recent reports on also support the notion [42], [43]. With the rapid enhancement in microbiome research, it has become evident that the manifestation of a viral infection in the human body is an outcome of a complex interplay between the host, the virus and the resident microbiota [44], [9].

Given the expected variations in the host microbiota across different regions in the world, the present study aimed at presenting the changes in the nasopharyngeal microbiota in COVID-19 positive individuals in an Indian context. This study design also focused on identifying the variations in the microbiota signatures between the symptomatic and asymptomatic individuals and present a set of microbes which might play a role in minimizing the symptoms of a respiratory infection. In addition, possibly for the first time, the study presents the change in the human nasopharyngeal microbiota over the course of the pandemic, which had varied presentation in all the three waves in India, largely caused by different SARS-CoV-2 variants (spanning the first two years). It revealed a perturbation in the microbial diversity and composition with a gradual drop in the abundance of opportunistic pathogens in the human respiratory tract across both the CP and CN groups. Overall, the findings are expected to enhance the general understanding of the human nasopharyngeal microbiota and aid in devising strategies to ward off other respiratory pathogens in the future.

## Conclusions

Most studies on the effect of COVID-19 infection on the human nasopharyngeal microbiota were conducted during the early period of the pandemic. Reports capturing the effect of the changing nature of the virus and/or interventions like vaccination, isolation (due to lockdown) on the host microbiota are largely missing. The current study examined the nasopharyngeal microbiota of COVID-19 positive as well as COVID-19 negative Indians and report the consistent and the varying patterns in the microbiota across the three COVID-19 waves, in terms of microbial diversity, taxonomy and inferred functions. In addition, variations between the microbiota samples collected from symptomatic and asymptomatic patients have also been reported. Whether these changes assist in COVID-19 disease onset and/or progression, would be interesting to explore in future.

## Methods

### Sample collection

This study was conducted for all three waves of COVID-19 in India, from March 2020 to February 2022 in accordance with the guidelines of the Indian Council of Medical Research (ICMR), Government of India and approved by the Institutional Ethics Committee of Centre for Cellular and Molecular Biology (CSIR-CCMB) [IEC-83/2020]. A total of 646 nasopharyngeal swab samples were collected from individual subjects, from different districts of Telangana state in India (data from 589 samples was used for further analysis after discarding 57 samples for various reasons including poor sequencing quality, depth, etc). The samples corresponded to the three COVID-19 waves in India and the time periods for swab collection for each wave were as follows: March 2020 to August 2020 for the first wave, April 2021 to July 2021 for the second, and January 2022 to February 2022 for the third. The swab samples were stored at -80°C until nucleic acid extraction, to ensure DNA quality. Subjects from each wave were divided into COVID-19 positive (CP) and COVID-19 negative (CN) groups. Among the CP group, subjects reporting severe symptoms and/or requiring hospitalization were further categorized as symptomatic COVID-19 positive (sCP). CP with no symptoms were denoted as asymptomatic COVID-19 positive (aCP). CN were further divided into asymptomatic (aCN) and symptomatic (sCN), i.e., COVID-19 negative subjects with presumably respiratory/other infections.

The samples used in this study were collected for SARS-CoV-2 diagnostics and genome sequencing at CSIR-CCMB, also an ICMR-approved COVID-19 testing center. The diagnosis of COVID-19 patients involved combining the results from the real-time reverse transcription-polymerase chain reaction (qRT-PCR) assay performed on nasopharyngeal swabs, at the BSL-2 facility at CSIR-CCMB.

### DNA isolation

DNA was extracted from the viral transport media (VTM) containing the nasopharyngeal swabs using the QIAamp^®^ DNA Microbiome Kit (QIAGEN, Hilden, Germany) according to the manufacturer’s instructions. All the extraction procedures were performed in the pre-PCR designated room in the BSL-2 facility. The yield of the extracted DNA was validated by Nanodrop Spectrophotometric (Thermofisher Scientific, Milan, Italy) reading at 260/280 as well as 260/230 and the quality was assessed by running an aliquot of the DNA in a 1% agarose gel.

### 16S rRNA gene amplification and sequencing

The first round of PCR was performed in a 20μL reaction mixture, with 50 ng of bacterial genomic DNA as the template. The primer pair targeting the V4 hypervariable region (515-F and 806-R) [45] of the 16S rRNA gene (see Supplementary Methods) was modified by adding the Illumina overhang forward and reverse adaptor sequences.

The component concentration of each reaction mix included 10μL of 2X EmeraldAmp^®^ GT PCR master mix (Takara Bio, Japan), 1μL (0.5μM) of each primer, and 7μL of Nuclease free water. Thermocycling was performed on a Bio-Rad T100 Thermal cycler and included an initial denaturation at 95°C for 3 min, followed by 35 cycles of 95°C for 30s, 55°C for 30s, and 72°C for 30s, followed by a final extension of 72°C for 5 min. Each PCR reaction mixture was loaded into a 2% agarose gel, stained with ethidium bromide to observe the amplification.

The second round of PCR involved attaching dual indices and Illumina sequencing barcodes to the purified 16S rRNA gene amplicons, which was performed using the Nextera XT Index kit as mentioned in the manufacturer’s protocol. The prepared metagenomic libraries were then quantified and normalized using Qubit dsDNA BR Assay before getting pooled in equimolar concentrations. Our constructed libraries were then sequenced (seven sequencing runs, Supplementary Table 9) in a paired-end mode (2 × 300) on the Illumina MiSeq sequencing platform using v3 600 cycles reagent.

### Bioinformatics analysis

The raw reads were processed using cutadapt (version 3.4) and were fed to the DADA2 pipeline (version 1.20) for ASV generation and subsequent taxonomic assignment. The alpha (observed ASVs and Shannon diversity) and beta diversity (Jaccard and Bray–Curtis distance) measures were computed using the Phyloseq package (version 1.26.1). Cytoscope (version 3.9.0) and NetShift ^24^ were utilized for constructing the correlational networks and for identifying the taxas driving the major shift in case of COVID-19 infection networks. The functional potential of the microbiota samples was predicted using PICRUSt2 [46]. The differentially abundant discriminating taxa and (inferred) functions were identified through discriminant analysis using the DESeq2 package (v 1.34.0). The detailed steps along with the parameters used for each of the bioinformatic analysis have been presented in the Supplementary Methods section.

## Declarations

### Ethics approval and consent to participate

This study was conducted for all three waves of COVID-19 in India, from March 2020 to February 2022 in accordance with the guidelines of the Indian Council of Medical Research (ICMR), Government of India and approved by the Institutional Ethics Committee of Centre for Cellular and Molecular Biology (CSIR-CCMB) [IEC-83/2020].

## Consent for Publication

All authors have read and approved the paper for submission.

## Availability of data and materials

All data generated and analyzed in this study are available within the article and as its Supplementary information files. The datasets supporting the conclusions of this article are available in the National Centre for Biotechnology Information-Sequence Read Archive (SRA) with the BioProject ID PRJNA902495.

## Author Contributions

MMH and SSM conceived the idea. ABS, AD, BV, DTS, KBT, MMH, SSM and TB created the study design. ABS, BV, KBT and DTS designed the sequencing protocol. VA and TN performed the wet lab, including sequencing experiments. WU, HK, MR, NKP, TB and JSK performed the bioinformatic analysis, ABS, AD, NKP, TB, VA and WU analyzed the results and wrote the manuscript. ABS and MMH managed the project and oversaw the overall progress. All authors read and approved the final version of the manuscript.

## Acknowledgements

Authors also acknowledge the guidance received from Dr. Rakesh Kumar Mishra during the course of the study. We are also grateful to Dr. Shreekant Verma and Dr. Dhiviya Vedagiri for their valuable suggestions during the study and to Venkatesh Madihalli for his cooperation and continued assistance.

## Supplementary Information

### List of Supplementary Figures

**Fig. S1:** Venn diagram representing the distribution of 8645 amplicon sequence variants (ASVs) across (**A**) four disease categories, and (**B**) three COVID-19 waves.

**Fig. S2:** Stacked bar-plot representing the distribution of the bacterial taxonomies across samples. Distribution of bacteria at (**A**) phylum and (**B**) family level in COVID-19 positive (CP) and COVID-19 negative (CN) samples across the three COVID-19 waves. Distribution of bacteria at (**C**) phylum and (**D**) family level between the four sample sub-groups: asymptomatic COVID-19 positive (aCP), symptomatic COVID-19 positive (sCP), apparently healthy controls who are COVID-19 negatives as well as asymptomatic (aCN), and COVID-19 negatives exhibiting symptoms (sCN).

**Fig. S3:** Differential abundance of bacterial phylum between the COVID-19 positive (CP) and COVID-19 negative (CN) samples. The log2fold change in the mean abundance (along with whiskers representing standard errors) of a bacterial phylum in CP with respect to CN is depicted in (**A**) all the analyzed samples (overall) as well as in (**B-D**) each of the three COVID-19 waves. Significantly different abundance (q-value < 0.05) is indicated with blue colour.

**Fig. S4:** Differential abundance of bacterial phylum between the COVID-19 positive (CP) and COVID-19 negative (CN) samples across the three COVID-19 waves. The log2fold change in the mean abundance (along with whiskers representing standard errors) of a bacterial phylum in the second wave with respect to the first wave in (**A**) CN samples, (**B**) CP samples; third wave with respect to the first wave in (**C**) CN samples, (**D**) CP samples; third wave with respect to the second wave in (**E**) CN samples, (**F**) CP samples. Significantly different abundance (q-value < 0.05) is indicated with blue colour.

**Fig. S5:** Differential abundance of bacterial phylum between the four sub-group of samples - asymptomatic COVID-19 positive (aCP), symptomatic COVID-19 positive (sCP), apparently healthy controls who are COVID-19 negatives as well as asymptomatic (aCN), and COVID-19 negatives exhibiting symptoms (sCN). The log2fold change in the mean abundance (along with whiskers representing standard errors) of a bacterial phylum in (**A**) aCP with respect to aCN, (**B**) sCN with respect to aCN, (**C**) sCP with respect to sCN, and (**D**) sCP with respect to aCP is depicted. Significantly different abundance (q-value < 0.05) is indicated with blue colour.

**Fig. S6:** Distribution of betweenness centralities of the nodes (microbes) in each of the analyzed microbial association network.

**Fig. S7:** The changes in community structure (community shuffling) between the microbial association networks corresponding to the first COVID-19 wave - ‘control’ and the second COVID-19 wave - ‘case’ networks. Nodes belonging to the ‘control’ and ‘case’ networks are plotted along the left half and right half of the circular frame. Same node (microbe) in the two network is connected by an edge for easy viewing of the community shuffling. Node labels are coloured (at random) based on sub-network/ community affiliations. Grayed out node labels indicate that the node does not interact directly with the common sub-network. The node sizes are proportional to the betweenness centrality measure of the node in the corresponding network.

**Fig. S8:** The changes in community structure (community shuffling) between the microbial association networks corresponding to the second COVID-19 wave - ‘control’ and the third COVID-19 wave - ‘case’ networks. Nodes belonging to the ‘control’ and ‘case’ networks are plotted along the left half and right half of the circular frame. Same node (microbe) in the two network is connected by an edge for easy viewing of the community shuffling. Node labels are coloured (at random) based on sub-network/ community affiliations. Grayed out node labels indicate that the node does not interact directly with the common sub-network. The node sizes are proportional to the betweenness centrality measure of the node in the corresponding network.

**Fig. S9:** The changes in community structure (community shuffling) between the microbial association networks corresponding to the samples from asymptomatic individuals - ‘control’ and the samples from symptomatic individuals - ‘case’ networks. Nodes belonging to the ‘control’ and ‘case’ networks are plotted along the left half and right half of the circular frame. Same node (microbe) in the two network is connected by an edge for easy viewing of the community shuffling. Node labels are coloured (at random) based on sub-network/ community affiliations. Grayed out node labels indicate that the node does not interact directly with the common sub-network. The node sizes are proportional to the betweenness centrality measure of the node in the corresponding network.

### List of Supplementary Tables

**Supplementary Table 1:** List of the amplicon sequence variants (ASVs) and genera demonstrating a consistent increase or decrease in abundance across the three COVID-19 waves. Significant changes having an adjusted p-value < 0.05 have been reported.

**Supplementary Table 2:** Discriminatory taxonomic groups identified at a ASV and genera level between the three COVID-19 waves among samples from the COVID-19 positive (CP) and COVID-19 negative (CN) groups.

**Supplementary Table 3:** Discriminatory taxonomic groups identified at a ASV and genera level between the four sample sub-groups based on symptoms and disease status - asymptomatic COVID-19 positive (aCP), symptomatic COVID-19 positive (sCP), apparently healthy controls who are COVID-19 negatives as well as asymptomatic (aCN), and COVID-19 negatives exhibiting symptoms (sCN).

**Supplementary Table 4:** Table of ‘Neighbor Shift (NESH) Score’ corresponding to Figure 5 and Figures S7-S9.

**Supplementary Table 5:** Discriminatory functional pathways identified between COVID-19 positive (CP) and COVID-19 negative (CN) samples across the three COVID-19 waves using inferred functional profiles obtained through PICRUSt2 analysis.

**Supplementary Table 6:** Discriminatory enzyme functional potentials identified between COVID-19 positive (CP) and COVID-19 negative (CN) samples across the three COVID-19 waves using inferred functional profiles obtained through PICRUSt2 analysis.

**Supplementary Table 7:** Discriminatory functional pathways identified between the four sample sub-groups based on symptoms and disease status - asymptomatic COVID-19 positive (aCP), symptomatic COVID-19 positive (sCP), apparently healthy controls who are COVID-19 negatives as well as asymptomatic (aCN), and COVID-19 negatives exhibiting symptoms (sCN), using inferred functional profiles obtained through PICRUSt2 analysis.

**Supplementary Table 8:** Discriminatory enzyme functional potentials identified between the four sample sub-groups based on symptoms and disease status - asymptomatic COVID-19 positive (aCP), symptomatic COVID-19 positive (sCP), apparently healthy controls who are COVID-19 negatives as well as asymptomatic (aCN), and COVID-19 negatives exhibiting symptoms (sCN), using inferred functional profiles obtained through PICRUSt2 analysis.

**Supplementary Table 9:** Statistics on the amplicon sequencing runs performed in this study.

