## Supplementary figures and images for "Composition of nasopharyngeal microbiota in individuals with SARS-CoV-2 infection across three COVID-19 waves in India"

### Supplementary Figure 1

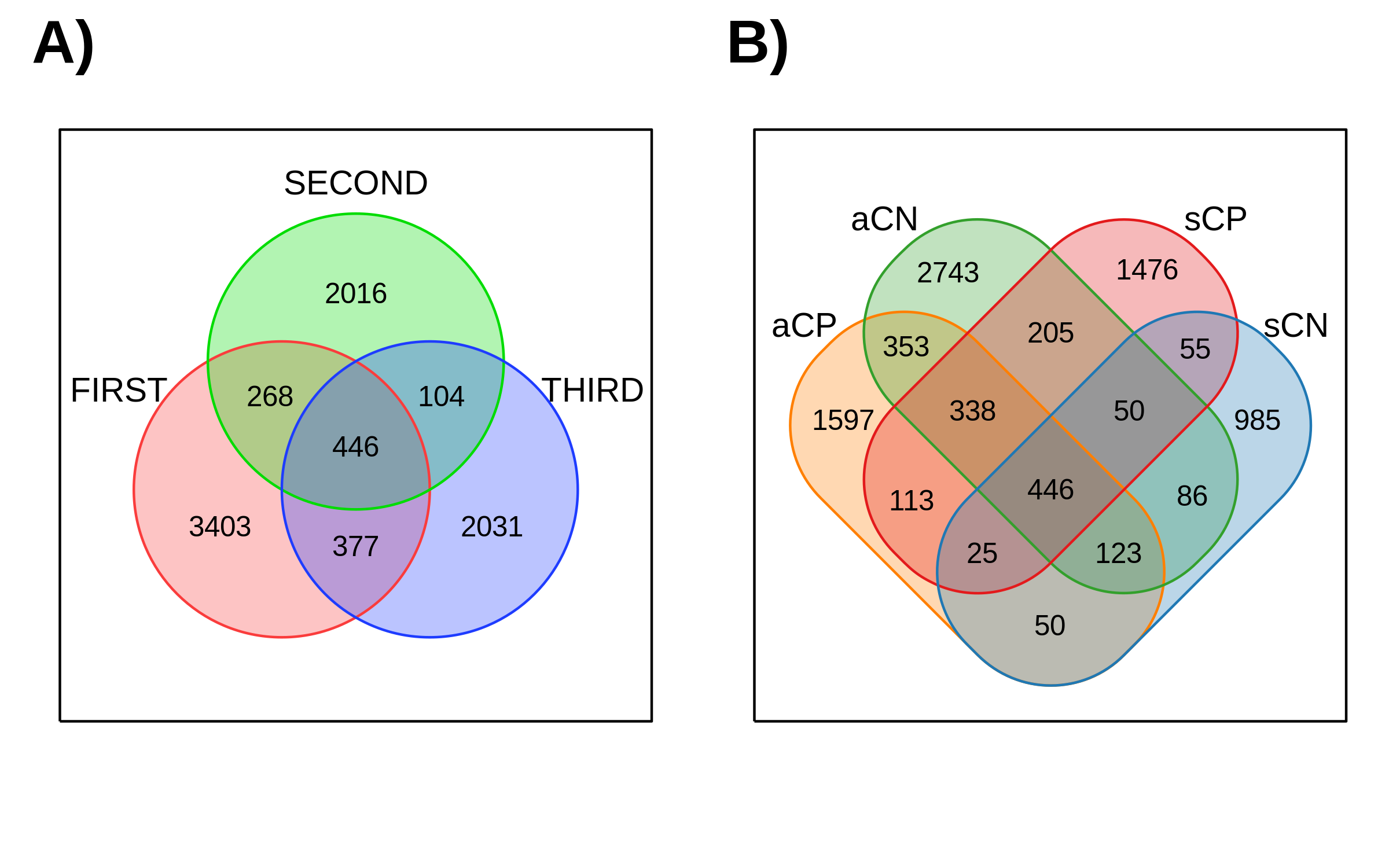

### Supplementary Figure 2

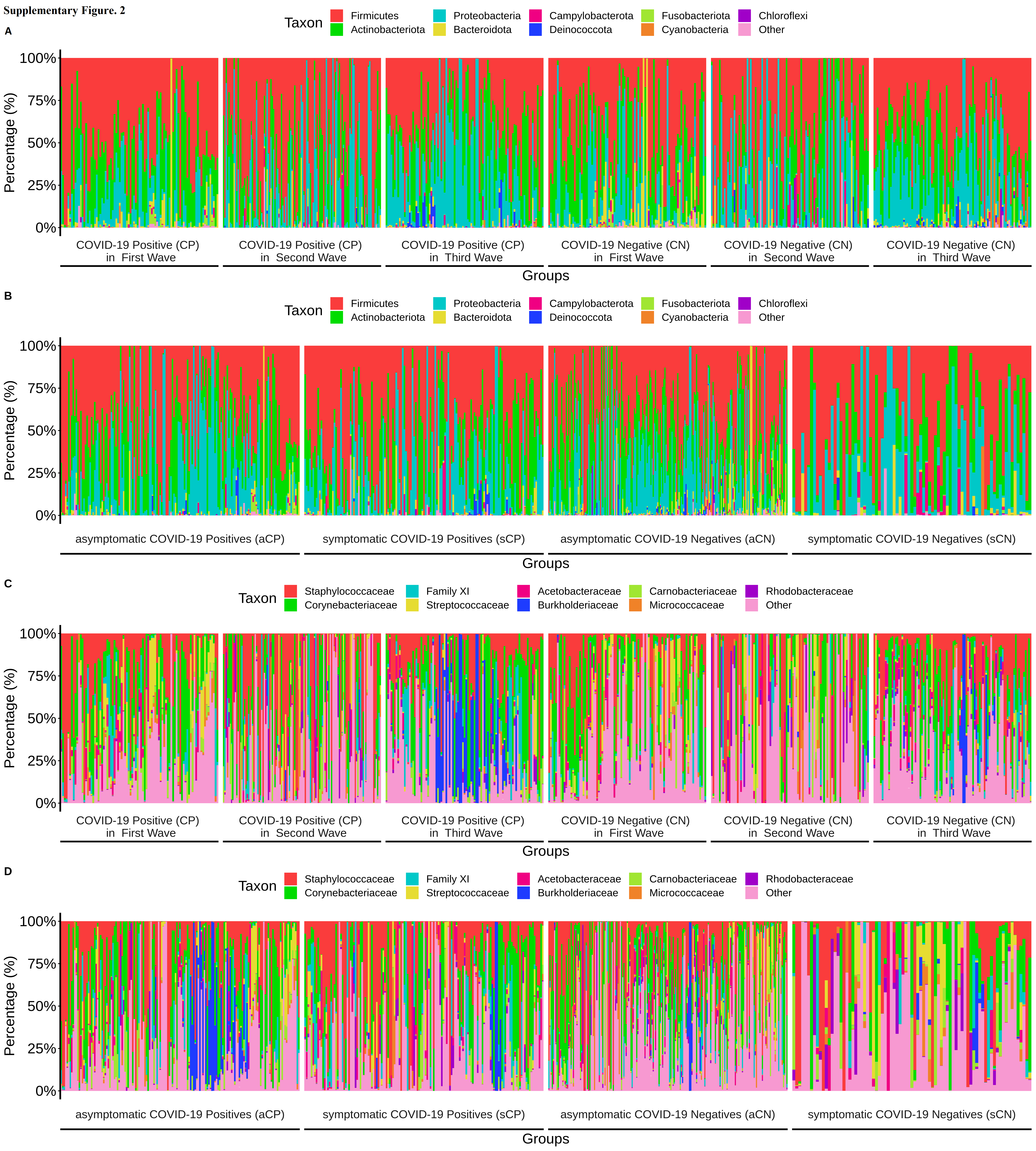

### Supplementary Figure 3

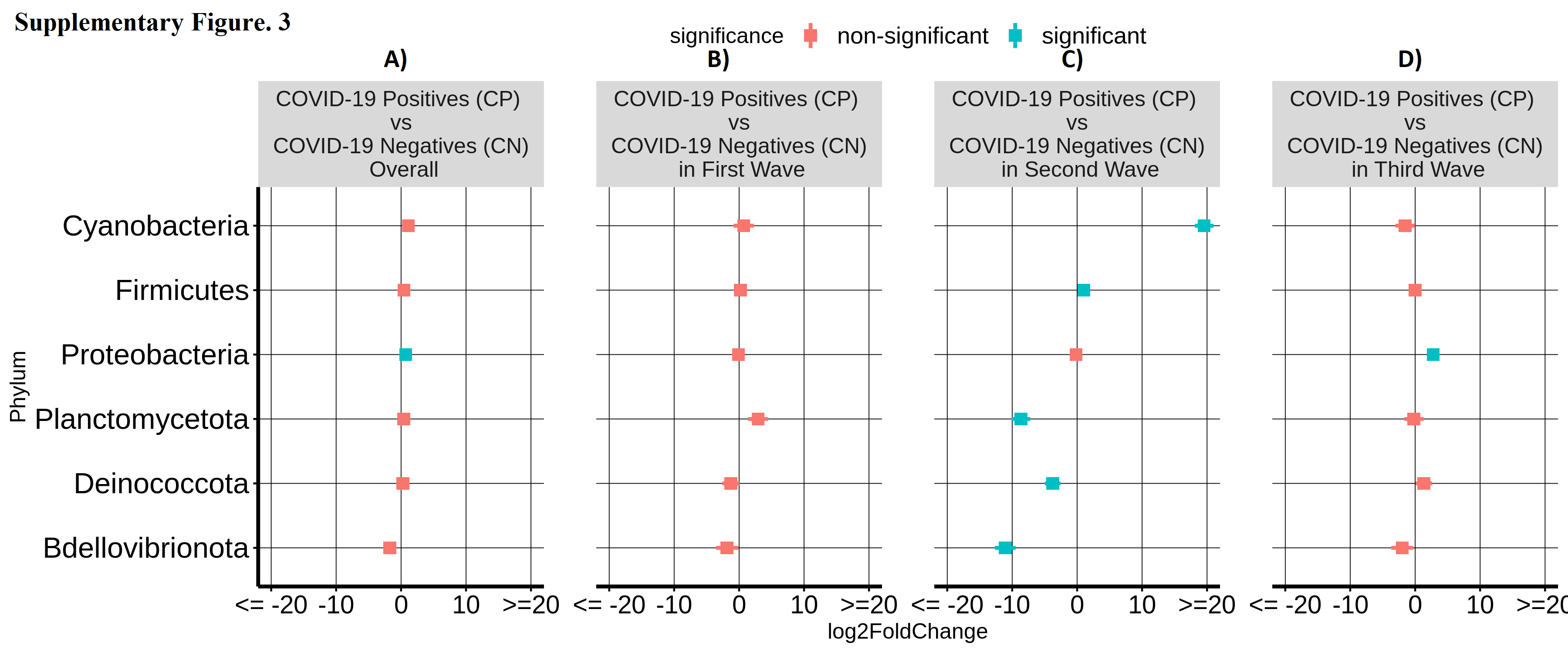

### Supplementary Figure 4

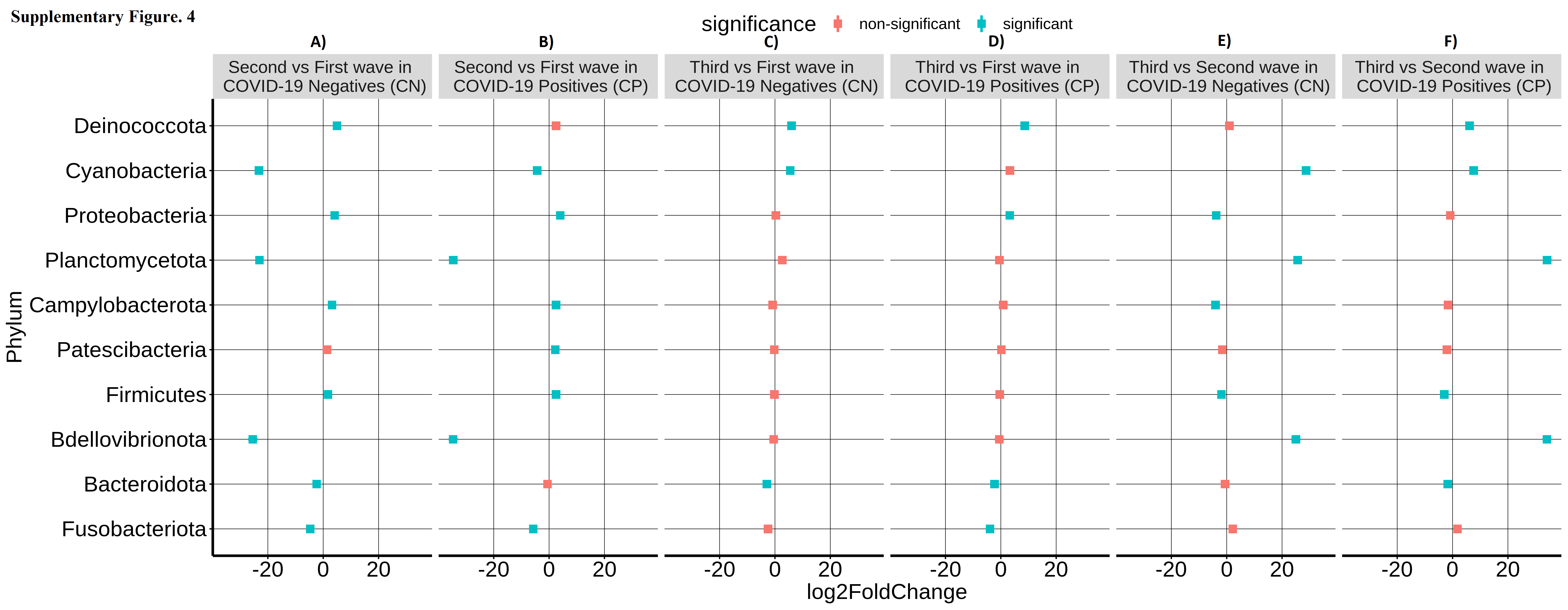

### Supplementary Figure 5

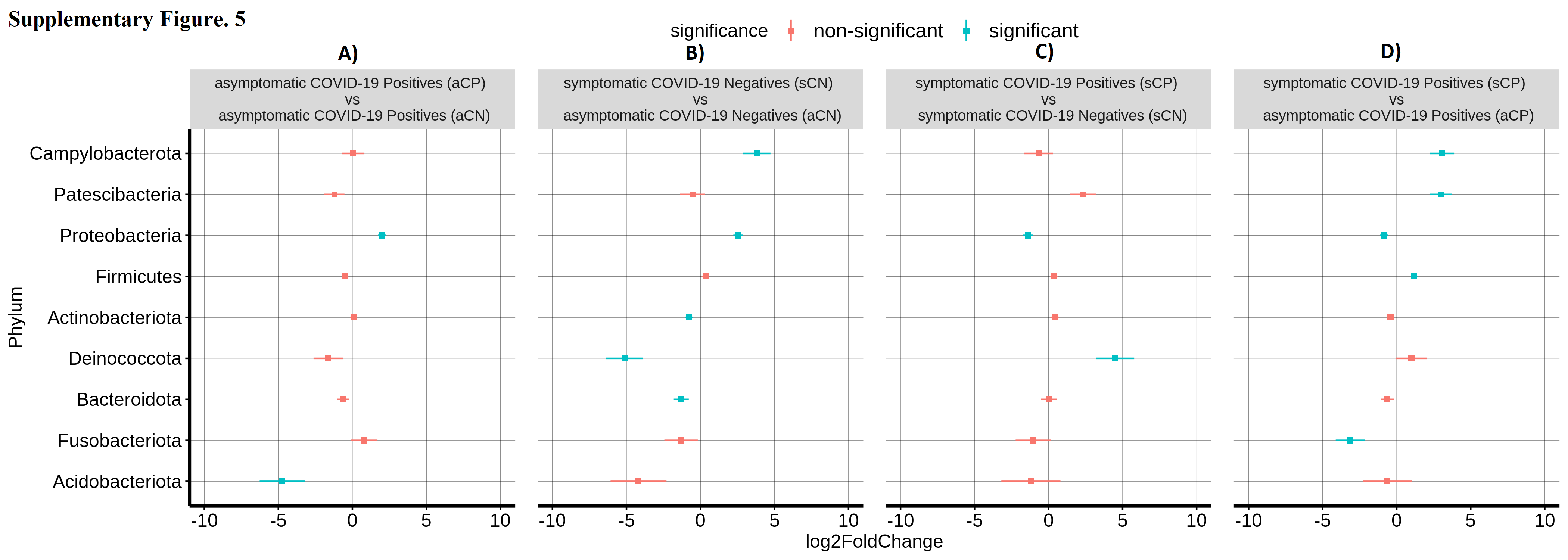

### Supplementary Figure 6

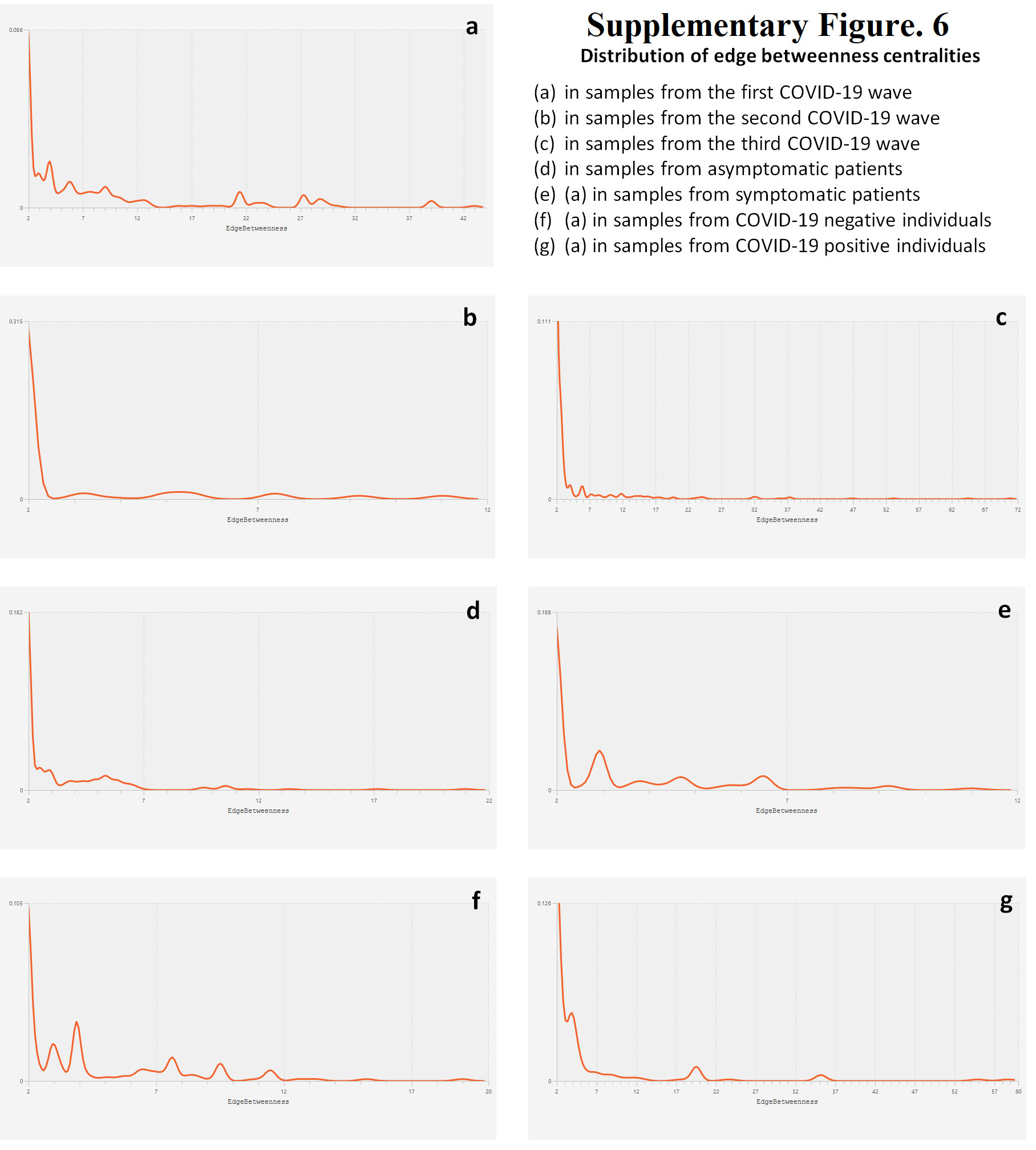

### Supplementary Figure 7

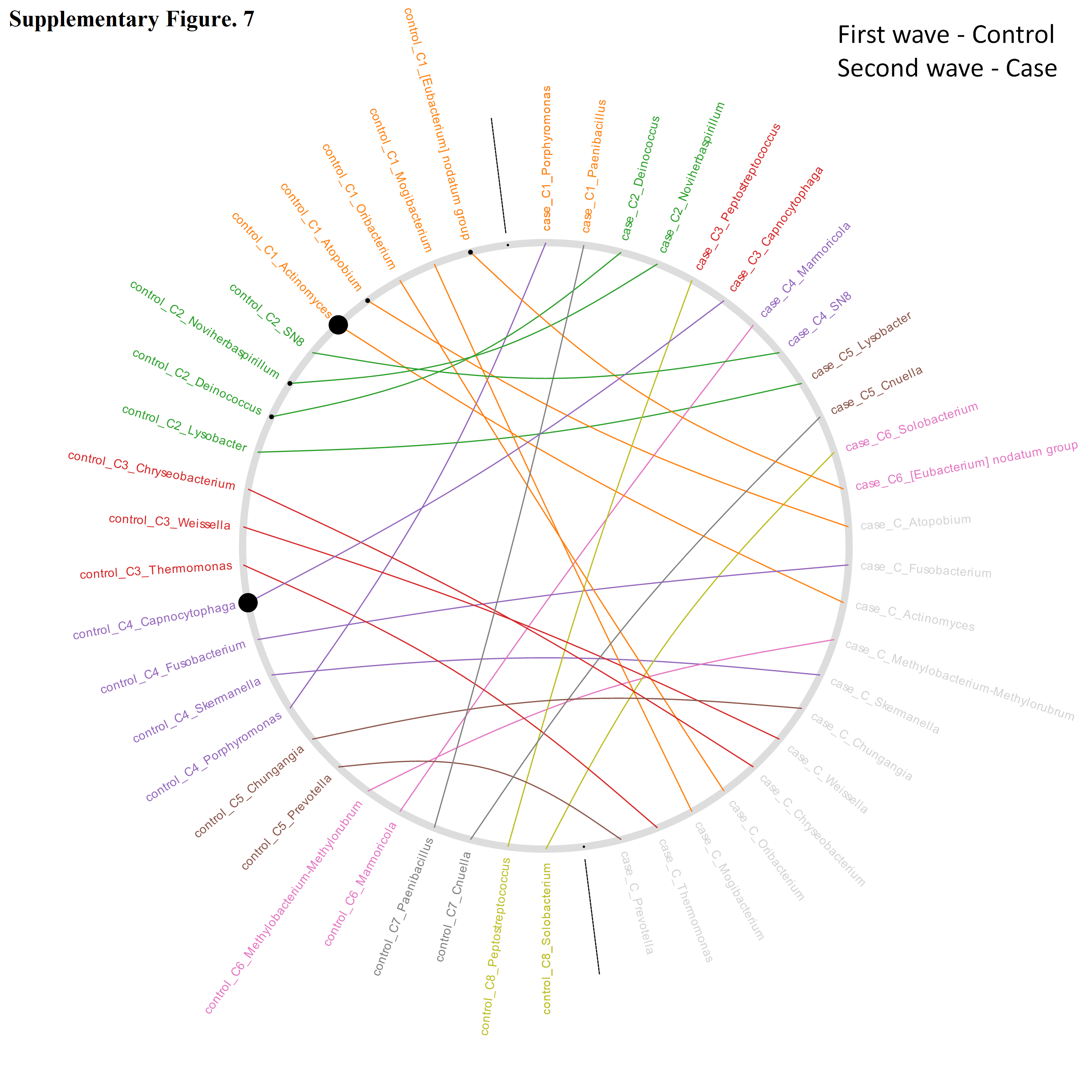

### Supplementary Figure 8

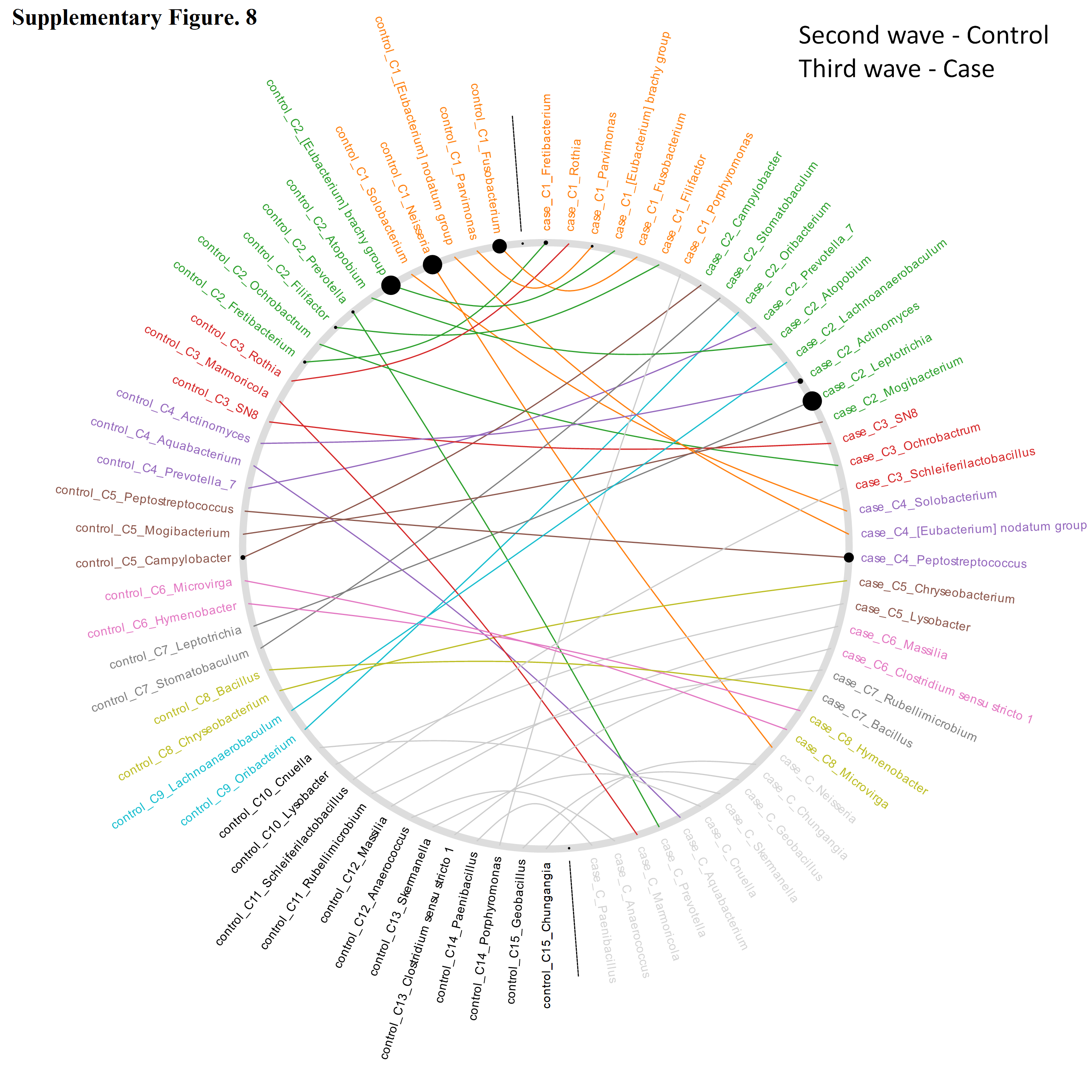

### Supplementary Figure 9

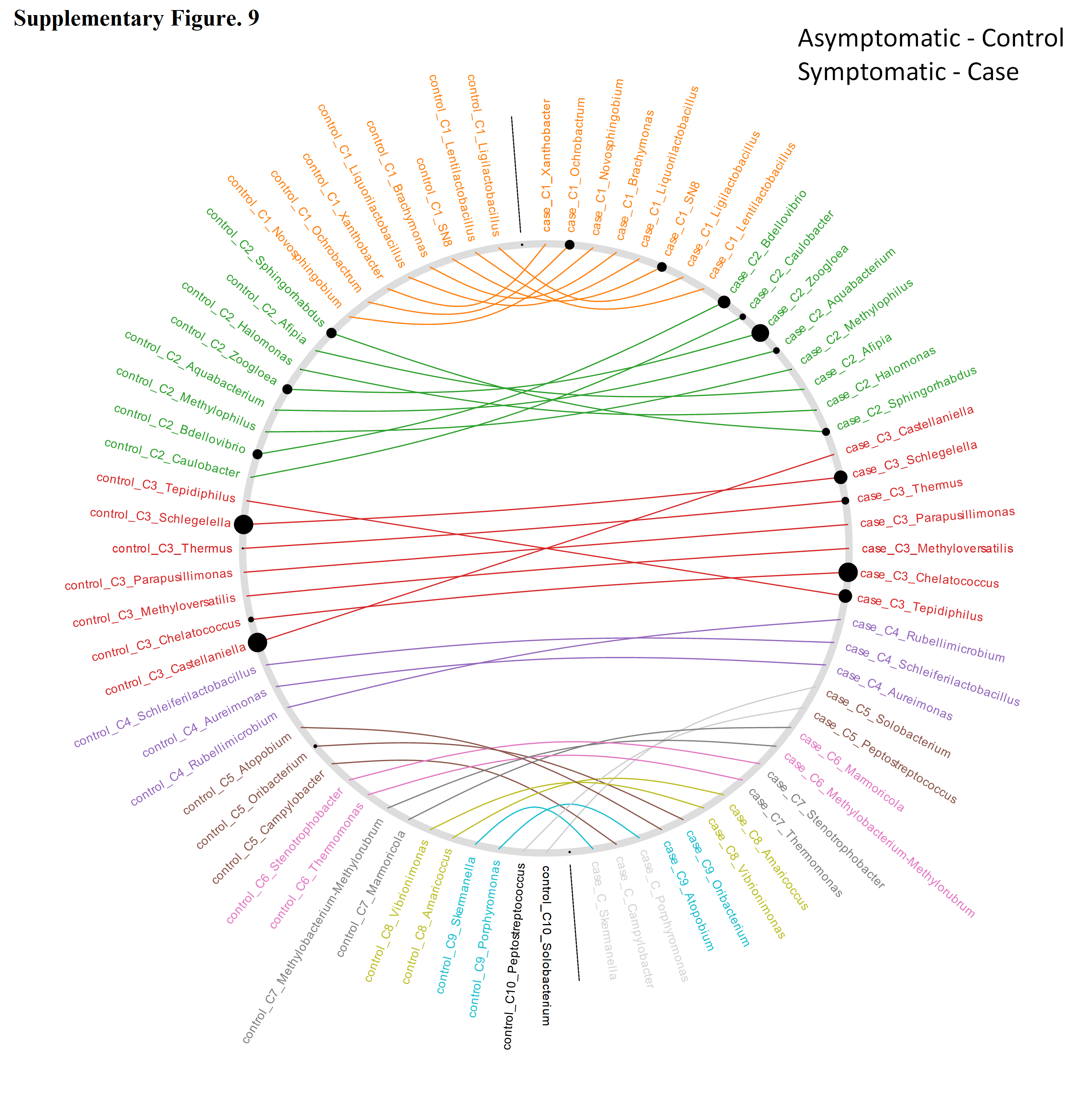
