## Supplementary MS for "Composition of nasopharyngeal microbiota in individuals with SARS-CoV-2 infection across three COVID-19 waves in India"

**Supplementary Results**

**Microbial taxa associated with symptomatic and asymptomatic subjects**

With respect to the microbiota samples from aCN subgroup, the signature of the aCP microbiota was characterized by a reduced abundance of Rickettsiaceae, Enterobacteriaceae (both belonging to phylum Proteobacteria), Moraxellaceae, Pseudomonadaceae (both belonging to phyla Pseudomonadota), Aerococcaceae, Thermaceae, and Micrococcaceae. At the same time, an enriched abundance of Burkholderiaceae, Xanthobacteraceae (both Proteobacteria) and Carnobacteriaceae were observed in aCP. Further, within the symptomatic subgroups, Thermaceae, Dysgonomonadaceae, Dermacoccaceae, and Alteromonadaceae were amongst the bacterial families with significant enrichments in sCP (w.r.t. sCN). In contrast, w.r.t. sCN, Rickettsiaceae was observed to be the most depleted family in sCP.

Notably, while the abundance of Rickettsiaceae decreased during COVID-19 infections, it has been reported to increase in other respiratory infections. Thus, it appears to be a distinguishing feature between COVID-19 and other respiratory infections. Similarly, enrichment of Thermaceae and Enterobacteriaceae, as well as a depleted abundance of Xanthobacteraceae appeared to be markers of a healthy nasopharyngeal microbiota since this trend was observed when aCN group was compared to both the sCN and aCP groups. *Kocuria* and *Liquorilactobacillus* were also linked to a healthy nasopharyngeal microbiota (Supplementary Table 4). Further, the opportunistic pathogenic genera *Stenotrophomonas* was noted to be depleted in a healthy respiratory tract microbiota.

**Variation in microbial networks between symptomatic and asymptomatic subjects**

The microbiome association network corresponding to the symptomatic condition (considering data from all three waves as well as both CP and CN groups combined) was found to be represented by a relatively lower number of microbial groups which also interacted less coherently amongst themselves (average number of neighbors), compared to the network corresponding to the asymptomatic condition (Table 4). From the NetShift constructed network (Supplementary Fig. 9), it was intriguing to note that the sub-network composition between the asymptomatic and symptomatic networks largely remained the same, with nodes belonging to the major four clusters showing no change in their cluster memberships. However, the betweenness centralities of some of these nodes changed between the symptomatic and the asymptomatic condition, indicating a change in the overall network architecture (Supplementary Fig. 9). Such changes were most evident in the clusters 2 and 3. Amongst other noticeable changes, the genera *Methyloversatilis* seemed to gain prominence (both an increased NESH score as well as a higher abundance) in the network representing the symptomatic condition (Supplementary Table 4). In contrast *Porphyromonas*, *Campylobacter*, and *Skermanella* were found to lose their importance in the third wave network due to a decrease in their abundance or that of their interacting partners (Supplementary Tables 4 and Supplementary Fig. 9).

**Association of inferred functions with microbiota samples during COVID-19**

Across the three COVID-19 waves, the metabolic pathway for methanogenesis from acetate (METH-ACETATE-PWY) was found to be significantly enriched in the microbiota corresponding to the CP samples Supplementary Table 5. In contrast, the pathways for anaerobic gondoate biosynthesis (PWY-7663), aerobic respiration I using cytochrome c (PWY-3781), cis-vaccenate biosynthesis (PWY-5973), Kdo transfer to lipid IVA III (PWY-6467), and chlorophyllide a biosynthesis I (CHLOROPHYLL-SYN) appeared to be depleted functions in the CP group as inferred using PICRUSt2. While IgA-specific metalloendopeptidase (EC:3.4.24.13) appeared to be the most enriched enzyme in the CP group, sedolisin (EC:3.4.21.100) was most enriched in the CN group (Supplementary Table 6).

In terms of the functional differences between the CP and CN groups in each of the three COVID-19 waves, while the minimum number of differentially abundant pathways were seen among the first wave samples, the quantum of change among the discriminating pathways was relatively smaller (log2foldchange < ±2) in the third wave samples (Supplementary Table 5). Further, there was no overlap between the metabolic pathways differentially expressed between CP and CN samples in the first two COVID-19 waves with those in the third COVID-19 wave. In the first two COVID-19 waves, spirilloxanthin and 2,2'-diketo-spirilloxanthin biosynthesis pathway (PWY-6581) and coumarins biosynthesis pathway (PWY-7398) were seen to be enriched in the CP group. In contrast, the depleted pathways in the CP group (w.r.t. CN group) included Kdo transfer to lipid IVA III pathway (PWY-6467), L-methionine biosynthesis III pathway (HSERMETANA-PWY), TCA cycle V and VI pathways (PWY-6969 and PWY-5913), aerobic respiration I pathway (PWY-3781), coenzyme B biosynthesis pathway (P241-PWY), and NAD biosynthesis I (from aspartate) pathway (PYRIDNUCSYN-PWY). Notably, of the pathways which were found to be depleted in CP samples from the first two COVID-19 waves, L-methionine biosynthesis III pathway (HSERMETANA-PWY) and TCA cycle V and VI pathways (PWY-6969 and PWY-5913) were found to be marginally enriched in the CP samples from the third COVID-19 wave.

Supplementary Tables 7 & 8 provide the list of functional pathways and enzymes which were found to be differentially enriched among the four disease categories. 4-hydroxyphenylacetate degradation (3-HYDROXYPHENYLACETATE-DEGRADATION-PWY), and 3-phenylpropanoate degradation (P281-PWY) appeared to be enriched in the microbiota of the CP samples when compared to CN irrespective of symptoms.

Irrespective of COVID-19 disease status, the nasopharyngeal microbiota of the symptomatic patients, i.e., sCP and sCN were seen to be enriched for the following functional pathways w.r.t. the asymptomatic patients – pyrimidine deoxyribonucleotide phosphorylation (PWY-7197), (aerobic) heme biosynthesis I (HEME-BIOSYNTHESIS-II), de novo biosynthesis II of adenosine (PWY-7220) and guanosine (PWY-7222) deoxyribonucleotides, and pyrimidine nucleobases salvage (PWY-7208). The sucrose degradation IV pathway (PWY-5384) was, however, noted to be enriched only in sCN (when compared to aCN) but depleted in sCP (when compared to aCP). In contrast, the superpathway of mycolyl-arabinogalactan-peptidoglycan complex biosynthesis (PWY-6404) was found to be enriched in CP (when compared to CN) only in the symptomatic sub-group. Conversely, the pathways for sucrose degradation IV (PWY-5384), O-antigen building blocks biosynthesis (OANTIGEN-PWY), and mixed acid fermentation (FERMENTATION-PWY) were noted to be enriched in the aCP sub-group while compared to aCN.

**Supplementary Discussion**

Irrespective of the disease status, the nasopharyngeal microbiota in Indian samples was found to be dominated by the phylum Firmicutes, Proteobacteria, and Actinobacteriota and the family Staphylococcaceae and Corynebacteriaceae. This is in line with the reports presented in earlier studies inspecting nasopharyngeal microbiota in other geographies [29], [50], [30]. However, we noted the proportion of Firmicutes being two to three times higher than Proteobacteria in the nasopharyngeal microbiota in Indian samples, especially in the CN group. This concurs with the observations reported in earlier studies from the US and Spain [50] [30], while is in contrast to those observed in Italians wherein Proteobacteria was found to be the most dominant organism [29]

.

In our study, the diversity of the nasopharyngeal microbiota was seen to be varied across the three waves (Fig. 2 and Supplementary Fig. 2). While we expected such changes given the behavioral changes adopted during COVID-19 pandemic, we were more intrigued to find that the microbes which were found to be distinguishing between CP and CN were also varied across the waves in most cases. For instance, while the abundances of Cyanobacteria, Firmicutes, Planctomycetota, Deinococcota and Bdellovibrionota were significantly different between the CP and CN groups in the second wave, Proteobacteria was significantly differing between the third wave CP and CN samples (Supplementary Fig. 5).

We noted an enrichment of the pathway for chitin derivatives degradation (PWY-6906) in sCP w.r.t. sCN. While similar observations were not made in the asymptomatic group (aCP w.r.t. aCN), given the recent interest in the role of chitin derivatives as anti-viral agents [48], [49] we thought this would be worth mentioning. This observation paves way for a deeper introspection into the microbiota’s potential in combating viral infection through the production of antiviral metabolites.

**Supplementary Methods**

**The primer pair targeting the V4 hypervariable region (515-F and 806-R)**

V4 515-F

**TCGTCGGCAGCGTCAGATGTGTATAAGAGACAG**GTGCCAGCMGCCGCGGTAA

V4 806-R	**GTCTCGTGGGCTCGGAGATGTGTATAAGAGACAG**GGACTACHVGGGTWTCTAAT

**Data QC and ASV generation:** A series of bioinformatics steps were followed to ensure usage of high-quality reads for the generation of the amplicon sequence variants (ASVs, i.e., sequence differing by as little as one nucleotide). The raw reads were processed using cutadapt (version 3.4) prior to their analysis through the DADA2 pipeline (version 1.20) for ASV generation and subsequent taxonomic assignment. Based on the quality of reads generated across the seven sequencing runs, the forward and reverse reads sequences were trimmed to 170 and 140 bases, respectively in the filterAndTrim step of DADA2. Furthermore, all read pairs with non-standard nucleotide bases, more than two ‘expected errors’ (maxEE), and lengths lower than 100 bases were discarded. Reads (and their pairs) which encountered atleast one base called with a quality score ≤ 8 (truncQ = 8) were also discarded. To account for the run-based biases, the ASVs were generated separately for each of the runs. The error models for each of the runs in the learnErrors step of DADA2 was performed using the parameters randomize=T and nbase=5e^+8^. To aid in faster computation, the denoising step (wherein the error model was employed for denoising) was preceded by a dereplication step (derepFastq). The initial sequences table generated for each of the seven sequencing runs (through the makeSequenceTable step of DADA2) were finally merged using mergeSequenceTables command of DADA2. Next, the removeBimeraDenovo step (method = “consensus”) was employed to remove ASVs originating from chimeric sequences. The retained ASVs were annotated using the assignTaxonomy function in DADA2 following the naïve Bayesian classifier method with the silva_nr_v132 database. Further, the species for these ASVs were assigned using addSpecies function with the silva_species_assignment_v132 database. As a final quality control step, ASVs with extremely low counts (n < 10 across all the sequenced samples) and irrelevant annotations (viz, Order classified as ‘chloroplast’ or ‘na’; Family classified as ‘mitochondria’ or ‘na’; Phylum classified as ‘uncharacterized’ or ‘na’) were also removed. The final microbiota data consisted of 8645 ASV belonging to 589 samples and were used for subsequent analysis.

**Alpha diversity:** To determine intra individual diversity, we calculated two alpha diversity measures: observed number of ASVs (observed ASVs) and Shannon diversity using the Phyloseq package (*phyloseq*, 1.26.1). Using generalized linear mixed models (GLMMs), we modeled alpha diversity according to COVID-19 status (CN; n=285, CP; n=304), wave (First; n=181, Second; n=217, Third; n=191), interaction between them (COVID-19*Wave), symptoms (Asymptomatic [aCP + aCN]; n=361, Symptomatic [sCP + sCN]; n=228), and sequencing depth to account for differential sequencing effort between samples, sequencing run and to control the random effect of batch processing of samples (*lme4*). In addition, we included the variables age group (25-29y; n=149, 30-34y; n=141, 35-39y; n=122, 40-44y; n=82, 45-50y; n=95), gender (female; n=276, male=313) and Ct-value as these factors could influence the microbiome. To facilitate model convergence, sequencing depth was scaled and to control the run effect we included run as a random factor variable. We used a log distribution for modeling the count data (observed ASVs) and a normal distribution for modeling the continuous data (Shannon diversity). Model selection was based on the information-theoretic (IT) approach using a second-order Akaike’s information criterion corrected for small sample sizes (AIC_C_) as an information criterion and Akaike weights (ω) to determine model support. For all GLMMs, we report both conditional and marginal coefficients of determination of each model (R^2^_GLMM(c)_, which explains the variance of both the fixed and random factors, and R^2^_GLMM(m)_, which explains the variance of the fixed factors only), which we calculated as the variance explained by the best model and the ΔAIC_C_. Finally, we performed Tukey’s HSD test to detect differences between each individual category (i.e., COVID-19*Wave) on the above performed GLMMs’ outcome.

**Beta diversity:** To assess the nasopharyngeal bacterial community composition between individuals, we calculated the Jaccard and Bray–Curtis distance matrices using the Phyloseq package (*phyloseq*, 1.26.1). Jaccard accounts for presence-absence of the taxa whereas Bray-Curtis in addition gives weight to taxa abundance. We tested for differences in microbial beta diversity for COVID-19 in different waves using the permutational multivariate analysis of variance (PERMANOVA) test with 999 permutations implemented in the *adonis* function of the vegan package (Vegan, 2.6.2). The fixed variables in our full model were the following: symptoms, age group, gender, sequencing run, Ct-value, and sequencing depth. We retained the sequencing run variable in our full model to statistically account for its model support. To understand whether COVID-19 reflects true shift in microbial community composition or differential spread (dispersion) of data points from their group centroid, we investigated the homogeneity of the variances of COVID-19 positive and negative category using the PERMDISP test implemented in the *betadisper* function of the vegan package. To visualize patterns of separation between different sample categories, Principal Coordinates Analysis (PCoA) plots were prepared based on the Bray–Curtis dissimilarity coefficient.

**Discriminant taxonomy analysis:** To identify the differential abundant discriminating taxa between different the nasopharyngeal samples of subjects with varying COVID-19 status, symptoms and across different waves; negative binomial Wald tests were performed using the DESeq2 package (v 1.34.0). For this analysis, only 2120 (out of 8645) ASVs which were present in at least two samples were considered. In order to obtain differentially abundant taxa between Covid-19 positive (CP) and Covid-19 negative (CN) after correcting for the wave effect (i.e., overall) Design-1 (design = ~ Wave + Covid) was adopted for DESeq2 analysis. Further, Design-2 (design = ~ Wave + Covid + Wave:Covid) was implemented to understand the individual and combined effects (interaction) of wave (Wave) and COVID-19 status (Covid) on the microbiota composition. In addition, to ascertain the effect of symptoms (sym_asym) in conjugation with COVID-19 status (Covid), Design-3 (design = ~ sym_asym + Covid + sym_asym:Covid) was used. For each of the designs, DESeq2 was performed with default parameters except for the size factor estimation. The size factor estimation was done using “poscounts”. The discriminating taxonomies were also analyzed for higher taxonomic levels. For this purpose, the ASVs were also aggregated at higher taxonomic levels to generate abundance tables corresponding to Phyla, Family and Genus and a protocol similar to the ASV level analysis was followed. To obtain the significant taxonomic groups across the comparisons log2FoldChange and lfcSE (log2FoldChange Standard Error) were plotted using the ggplot (v 3.3.5) package in R environment (R version 4.1.1, R Development Core Team, [2011](https://onlinelibrary.wiley.com/doi/10.1111/1755-0998.13215#men13215-bib-0038)) software.

**Network construction and identification of driver taxa:** Correlational networks were constructed from the microbial abundance data to understand associations among the interacting taxa. All analyses were performed at the Genus level to have a dataset with a reasonable number of nodes for a good display and meaningful interpretation thereof. For generating the networks, a Pearson correlation coefficient cut-off of ±0.5 was used. The networks were generated using Cytoscope (v 3.9.0) and analyzed using its ‘analyze network’ module. Further, the taxa driving the major shift in case of COVID-19 infections were identified using NetShift [47]. The tool is hosted on <https://web.rniapps.net/netshift/> and uses ‘Neighbor Shift (NESH) Score’ to identify key taxonomic groups which are more likely to drive changes from a control/ healthy, to a case/disease state.

**Discriminant functional analysis:** The functional potential of the microbiota samples was predicted using PICRUSt2 [46]. Next discriminant analysis on the inferred functions (enzymes and metabolic pathways) were performed using the DESeq2 package (v 1.34.0). A similar design as elucidated for the discriminant taxonomy analysis was adopted in this case.
